# Direct activation of an innate immune system in bacteria by a viral capsid protein

**DOI:** 10.1101/2022.05.30.493996

**Authors:** Tong Zhang, Hedvig Tamman, Kyo Coppieters’t Wallant, Tatsuaki Kurata, Michele LeRoux, Sriram Srikant, Tetiana Brodiazhenko, Albinas Cepauskas, Ariel Talavera, Chloe Martens, Gemma C. Atkinson, Vasili Hauryliuk, Abel Garcia-Pino, Michael T. Laub

## Abstract

Bacteria have evolved sophisticated and diverse immunity mechanisms to protect themselves against a nearly constant onslaught of bacteriophages^1–3^. Similar to how eukaryotic innate immune systems sense foreign invaders through pathogen-associated molecular patterns (PAMPs)^4^, many bacterial immune systems that respond to bacteriophage infection require a phage-specific trigger to be activated. However, the identities of such triggers and the mechanistic basis of sensing remain almost completely unknown. Here, we discover and investigate the anti-phage function of a fused toxin-antitoxin (TA) system called CapRel^SJ46^ that protects *E. coli* against diverse phages. Through genetic, biochemical, and structural analysis, we demonstrate that the C-terminal domain of CapRel^SJ46^ regulates the toxic N-terminal region, serving as both an antitoxin element and a phage-infection sensor. Following infection by certain phages, the newly synthesized major capsid protein binds directly to the C-terminal domain of CapRel^SJ46^ to relieve autoinhibition, enabling the toxin domain to then pyrophosphorylate tRNAs, which blocks translation to restrict viral infection. Collectively, our results reveal the molecular mechanism by which a bacterial immune system directly senses a conserved, essential component of phages, suggesting a PAMP-like sensing model for TA-mediated innate immunity in bacteria. We provide evidence that CapRels and their phage-encoded triggers are engaged in a Red Queen conflict^5^, revealing a new front in the intense coevolutionary battle being waged by phage and bacteria. With capsid proteins of some eukaryotic viruses known to stimulate innate immune signaling in mammalian hosts^6–10^, our results now reveal an ancient, deeply conserved facet of immunity.

---

Innate immunity in eukaryotes relies on pattern recognition receptors that directly sense pathogen-associated molecular patterns (PAMPs), which are conserved molecules like bacterial lipopolysaccharide and flagellin, or viral RNA or DNA^4^. These innate immune signaling pathways must remain silent prior to infection, but be poised for rapid activation to defend against foreign invaders. Bacteria also encode innate immune systems to protect themselves against diverse invading bacteriophages, but how they sense infection is poorly understood. One exception is restriction-modification (RM) systems, which are effectively in constant surveillance mode, using DNA methylation to distinguish self from non-self. Similarly, for CRISPR-Cas systems, the adaptive immune system of some bacteria, guide RNAs enable a cell to specifically target foreign DNA. Dozens of new bacterial defense systems have been discovered in recent years^11–15^, but unlike RM and CRISPR-Cas, many of them must be specifically activated upon phage infection. This is particularly critical for abortive infection (Abi) systems in which a defense system uses a lethal effector to kill an infected cell and prevent propagation of the virus through a population^16^. The phage-encoded triggers for such bacterial immunity mechanisms are largely unknown.

Toxin-antitoxin (TA) systems are prevalent genetic elements in bacteria that are emerging as key components of anti-phage innate immunity^13,14,17,18^, often serving as abortive infection modules that kill infected cells to prevent spread of phages through a population. How TA systems sense and respond to phage infection remains poorly understood. For the *toxIN* system, toxin (ToxN) activation relies on efficient, phage-induced shutoff of host transcription coupled to the intrinsically fast turnover of the antitoxin *toxI*^19–21^. However, *toxI* is an RNA, whereas most TA systems feature a protein antitoxin. For systems with a protein antitoxin, the mechanism of activation is often assumed to arise through antitoxin degradation. Although protein antitoxins are often more proteolytically unstable than their cognate toxins, their turnover may not be fast enough to enable toxin activation on the time-scale of a phage infection^22^, suggesting the existence of alternative mechanisms for TA activation. Bacterial retrons function as tripartite TA systems and can be activated by overexpressing various prophage genes^23^, but whether these activators function as such during phage infection is unknown.

## CapRel^SJ46^ is a fused, anti-phage toxin-antitoxin system

To investigate the molecular basis of phage-induced activation of bacterial immunity, we focused here on toxSAS TA systems, which feature toxins homologous to bacterial small alarmone synthetases (SAS) that pyrophosphorylate purine nucleotides^24^. While most housekeeping alarmone synthetases produce the growth regulator (p)ppGpp^25,26^, toxSAS toxins can synthesize (p)ppApp to deplete ATP^24,27^ or pyrophosphorylate tRNAs to inhibit translation^24,28^. Their cognate antitoxins can either bind and neutralize the toxin or act as hydrolases to reverse toxin-catalyzed pyrophosphorylation^24,28^. One subfamily of translation-inhibiting toxSAS is called CapRel based on their prevalence in Cyanobacteria, Actinobacteria, and Proteobacteria and sequence similarity to the (p)ppGpp synthetase/hydrolase Rel. This subfamily includes a number of representatives that are, in contrast to canonical bicistronic TA systems, encoded by a single open reading frame, with an N-terminal domain homologous to toxSAS toxins and a C-terminal domain homologous to the corresponding antitoxins^29^ (Fig. 1a and S1a).

**Figure 1.**
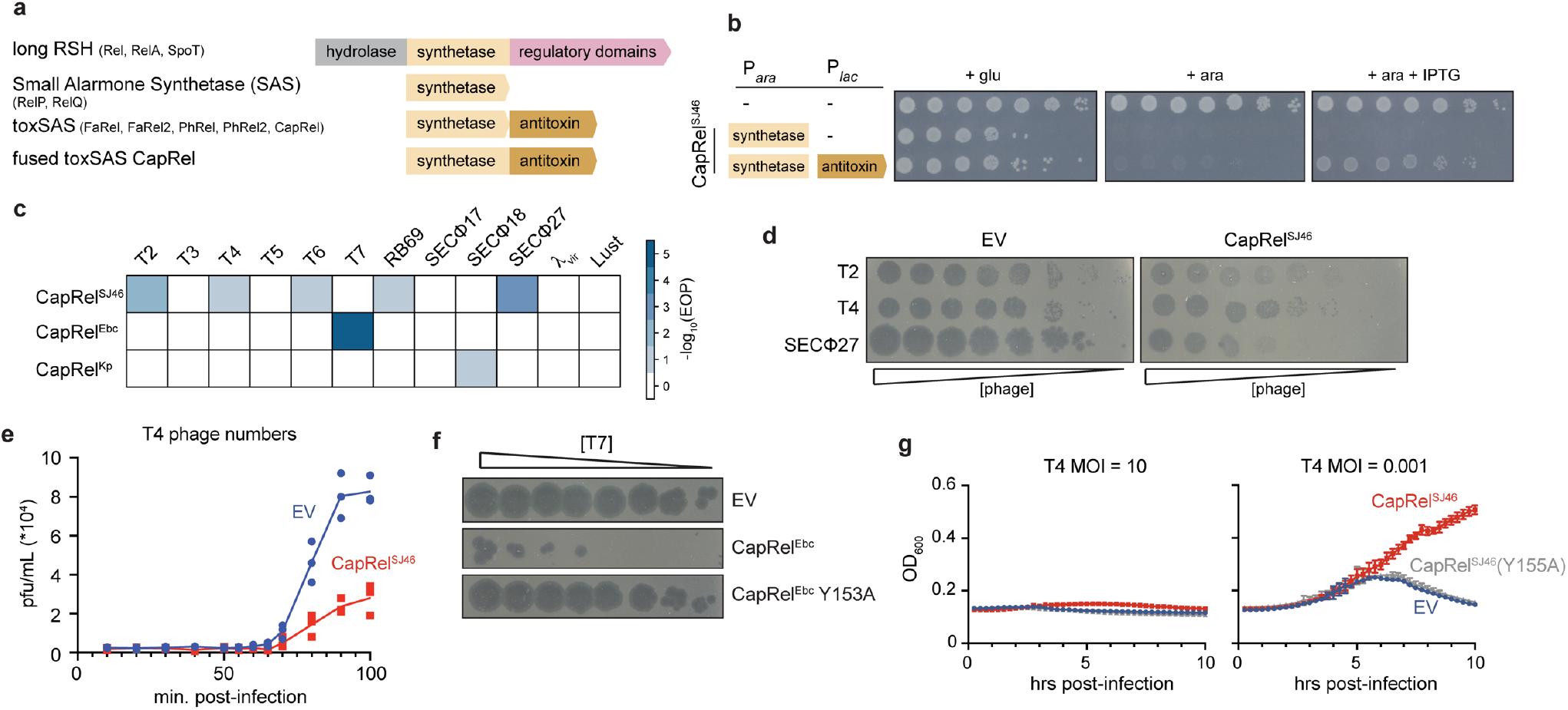
Fused CapRel homologs are toxin-antitoxin systems that can provide *E. coli* with robust defense against phages. (**a**) Domain organization of long RSH (RelA-SpoT Homologs), SAS (Small Alarmone Synthetases), toxSAS, and the fused subclass of toxSAS TA systems including CapRel^SJ46^. (**b**) Cell viability assessed by serial dilutions for strains expressing the N-terminal toxin domain of CapRel^SJ46^ alone or with the C-terminal antitoxin domain. (**c**) Efficiency of plaquing (EOP) data for the phages indicated when infecting cells producing CapRel^SJ46^, CapRel^Ebc^, or CapRel^Kp^. (**d**) Serial dilutions of the phages indicated spotted on lawns of cells harboring CapRel^SJ46^ or an empty vector (EV). (**e**) One-step growth curve measuring plaque forming units (pfu/mL) during the first round of infection by T4 of cells harboring CapRel^SJ46^ or an empty vector. (**f**) Serial dilutions of T7 phage spotted on lawns of cells harboring CapRel^Ebc^, CapRel^Ebc^(Y153A), or an empty vector. (**g**) Growth of cells producing CapRel^SJ46^ or CapRel^SJ46^(Y155A), or harboring an empty vector, following infection with T4 at a multiplicity of infection (MOI) of 10 (*left*) or 0.001 (*right*).

We selected a fused CapRel encoded by the *Salmonella* phage SJ46 and also encoded (with 100% amino acid sequence identity) in prophages of several *E. coli* strains (Fig. S1b). The toxin and antitoxin-like regions of CapRel^SJ46^ are related to the PhRel toxSAS toxin and its antitoxin ATphRel, respectively, from the mycobacterial temperate phage Phrann^30^ (Fig. S1a). This Phrann-encoded system can inhibit superinfection by other temperate mycophages^30^, although the molecular basis of PhRel activation is not known. To test if CapRel^SJ46^ is a fused TA system, we cloned the N-terminal region containing the conserved alarmone synthetase domain and the C-terminal region containing the putative antitoxin domain under the control of separate inducible promoters. Expression of the N-terminal fragment alone was toxic, and its toxicity was rescued *in trans* by co-expression with the C-terminal fragment (Fig. 1b), suggesting that CapRel^SJ46^ is a fused TA system.

To determine whether fused CapRels can defend against phages, we transformed *E. coli* MG1655 with three different systems expressed from their native promoters on low copy-number plasmids, and then tested whether each conferred protection against a panel of 12 diverse coliphages. In addition to CapRel^SJ46^, we also tested CapRel^Ebc^ from *Enterobacter chengduensis* and CapRel^Kp^ from *Klebsiella pneumoniae* (Fig. 1c and S1b-c). CapRel^SJ46^ decreased the efficiency of plaquing (EOP) for T2, T4, T6, RB69, and SECΦ27 by 10-1000-fold (Fig. 1c-d), indicating that this system provides strong protection against phages. T4 phage formed smaller plaques when plated onto CapRel^SJ46^-containing cells, and one-step growth curves confirmed that CapRel^SJ46^ reduces the burst size of T4 by ∼70% (Fig. 1e). CapRel^Ebc^ protected strongly against T7 and CapRel^Kp^ protected, albeit less efficiently, against SECΦ18 (Fig. 1f and S1c).

Next, we tested whether CapRel^SJ46^ provides direct immunity or functions through abortive infection in which an infected cell dies, but prevents the production of mature virions, thereby sparing uninfected cells in a population. To this end, we infected cells containing CapRel^SJ46^ with T4 at a multiplicity of infection (MOI) of either 10 or 0.001, and found that defense only manifested at the low MOI indicating that CapRel^SJ46^ likely functions through abortive infection (Fig. 1g). Phage protection by CapRel^SJ46^ depended on the predicted enzymatic activity of the N-terminal synthetase domain, as substituting the conserved tyrosine (Y155A) in the G-loop that is critical for substrate binding abolished phage protection^31^ (Fig. 1g). A similar catalysis-compromising substitution Y153A in CapRel^Ebc^ also abolished phage protection (Fig. 1f). Collectively, our results established that fused CapRels can provide anti-phage defense, with variable phage specificity.

To understand what determines the specificity of phage protection by fused CapRels, we compared CapRel^SJ46^ and CapRel^Ebc^. These two proteins share 70% amino acid identity overall, but harbor significant differences in their C-terminal regions, which are only 47% identical (Fig. 2a). In addition, this region is the least conserved when we compared a more diverse set of fused CapRel homologs (Fig. S2a). Because CapRel^SJ46^ and CapRel^Ebc^ protected against different phages, we made a chimera in which the C-terminal region of CapRel^SJ46^ was replaced by the corresponding region of CapRel^Ebc^. This chimeric CapRel no longer protected against SECΦ27 and gained protection against T7 (Fig. 2b), manifesting as decreased EOP and smaller plaques. This result indicates that the C-terminal region of CapRel is critical to phage specificity.

**Figure 2.**
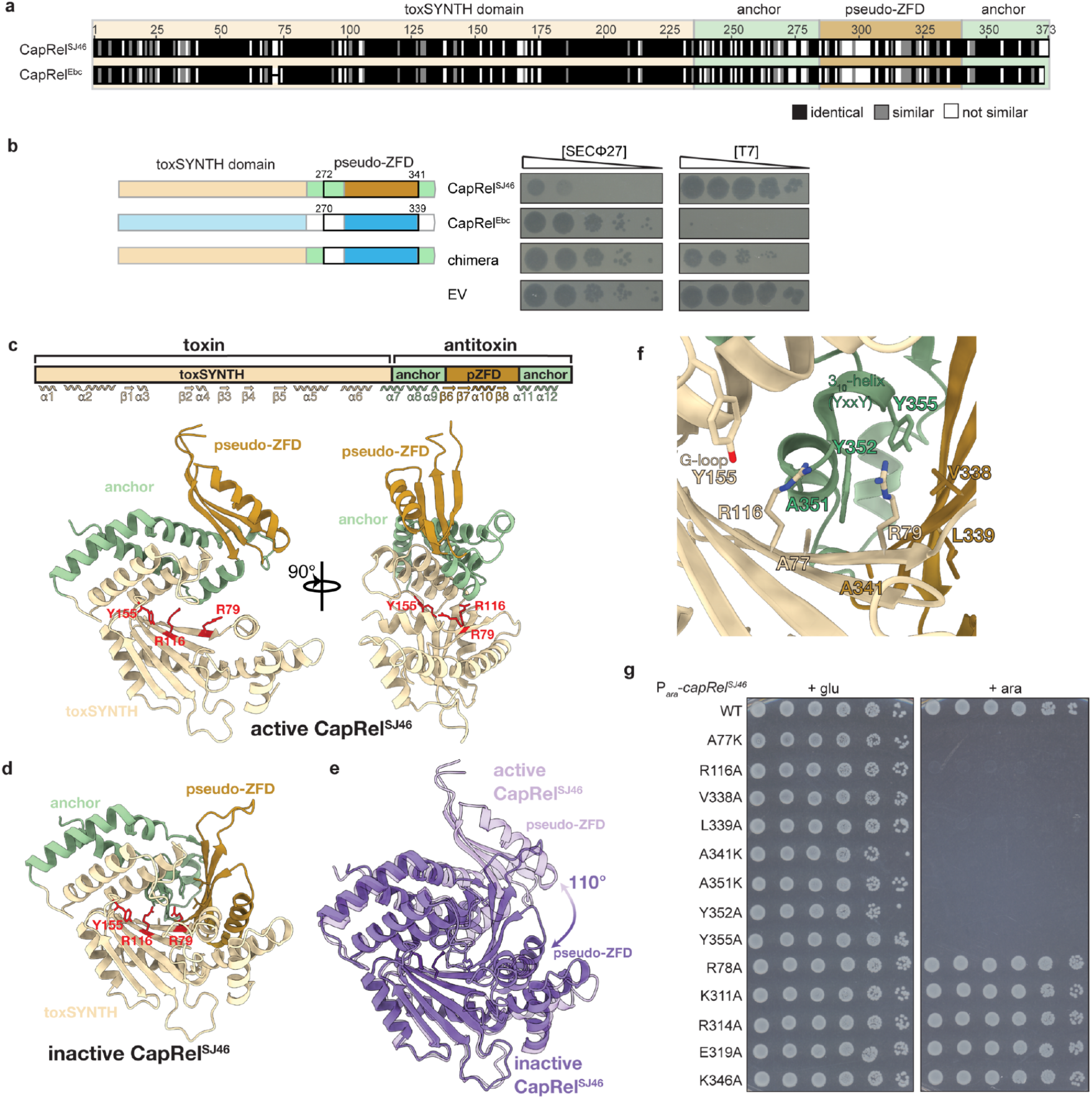
The pseudo-zinc finger antitoxin domain of CapRel confers phage specificity. (**a**) Sequence alignment of CapRel^SJ46^ and CapRel^Ebc^, with the more variable pseudo-zinc finger domain (pseudo-ZFD) labeled. (**b**) Serial dilutions of the phages indicated spotted on lawns of cells harboring the CapRel constructs indicated and diagrammed (*left*). (**c**) Cartoon representation of the structure of CapRel^SJ46^ with active site G-loop Y155 and the ATP-coordination residues R79 and R116 highlighted in red. Structural elements (toxSYNTH, pseudo-ZFD and the anchors) are coloured as in (**a**). (**d**) Closed conformation of CapRel^SJ46^ predicted by AlphaFold and coloured as (c). (**e**) Superposition of the active (open, light purple) and inactive (closed, dark purple) states of CapRel^SJ46^ as observed in the crystal structure and predicted by AlphaFold. (**f**) Details of the autoinhibited active site of CapRel^SJ46^ in the closed state. In this conformation, the YxxY neutralization motif of the pseudo-ZFD blocks the adenine coordination site, preventing catalysis. (**g**) Serial dilutions of cells expressing the indicated variant of CapRel^SJ46^ from an arabinose-inducible promoter on media containing glucose (*left*) or arabinose (*right*).

### Structural analysis of CapRel^SJ46^ reveals an autoinhibited and an active state

To further understand the mechanistic basis of anti-phage defense by CapRel^SJ46^, we solved a crystal structure to 2.3 Å resolution (Fig. 2c and Table S1). CapRel^SJ46^ contains a conserved, N-terminal nucleotide pyrophosphokinase domain present in alarmone synthetases and tRNA-pyrophospho-transferase enzymes, that mediates toxicity (toxSYNTH). The smaller C-terminal antitoxin domain consists of a central antiparallel three-stranded β-sheet with an α-helix connecting β-strands β7 and β8 (Fig. 2c, S2b-c). The antitoxin domain is topologically analogous to the classical Zn-finger domain (ZFD), but is lacking the conserved cysteines (Fig. S2d); we refer to this domain as a pseudo-ZFD. The pseudo-ZFD is connected to the toxSYNTH domain via α-helices α7, α8, and α9 and has a C-terminal α-helical extension that anchors the domain to α8 and α9 (Fig. S2b). In this structure the ATP donor nucleotide binding pocket and the conserved G-loop Y155 of toxSYNTH are exposed (Fig. 2c), indicating that this likely represents the active, toxic conformation of CapRel^SJ46^.

To explore the conformational dynamics of the enzyme, we used AlphaFold^32^ to predict possible alternative structures of CapRel^SJ46^. In addition to predicting the open conformation observed in the crystal structure (Fig. S2e), AlphaFold also predicted a closed conformational state in which the C-terminal domain folds back 110° onto the toxSYNTH central β-sheet and blocks the ATP-binding site (Fig. 2d). Comparison of the two states suggests that a conserved YxxY motif (Fig. S2a) located in the hinge connecting the two C-terminal α-helices in the open state morphs into a short 3_10_-helix in the closed state (Fig. 2e). This 3_10_-helix projects into the toxSYNTH active site and intercalates between β1 R79 and β2 R116 to block the adenine coordination site (Fig. 2e-f).

We hypothesized that this closed-to-open switch underlies the activation of CapRel^SJ46^, with the docking of the pseudo-ZFD onto toxSYNTH precluding substrate binding in the absence of phage infection (Fig. 2f). To test this hypothesis, we made single substitutions to the YxxY motif (Y352A and Y355A) and residues from the predicted interface that serves as a scaffold to orient and stabilize the 3_10_-helix (A77K, R116A, V338A, L339A, A341K, A351K), which are highly conserved among diverse CapRel homologs (Fig. S2a and S2f). Whereas wild-type CapRel^SJ46^ was not toxic when expressed in cells, each of the substitutions predicted to disrupt the intra-molecular recognition interface, on either the N- or C-terminal domain, rendered CapRel^SJ46^ toxic (Fig. 2g). These substitutions likely lead to constitutive activation of CapRel^SJ46^ by disrupting an autoinhibited state. As a control, we showed that substitutions in different structural elements of the pseudo-ZFD but not pointing toward the interface did not lead to constitutive activation (Fig. 2g). Collectively, our results indicate that the pseudo-ZFD docks onto the ATP-binding site of CapRel^SJ46^ to prevent switching to the open state captured in our crystal structure. Conservation of the YxxY motif and the interface residues suggest that this auto-inhibitory regulation is likely conserved in other CapRels.

### Fused CapRel^SJ46^ is activated by the major capsid protein of SECΦ27

Because full-length, wild-type CapRel^SJ46^ was not toxic when expressed in the absence of phage infection, we inferred that it must somehow be activated by phage. The toxins of some TA systems are activated by the degradation of the more labile antitoxin^19,33,34^. To test whether the C-terminal antitoxin of CapRel^SJ46^ is proteolytically cleaved off and degraded upon phage infection, we N-terminally tagged CapRel^SJ46^ and first verified that the tagged protein still defends against phage (Fig. S3a). We then tracked the size of CapRel^SJ46^ by immunoblotting following infection with SECΦ27. The overall protein levels of CapRel^SJ46^ remained constant and we observed only the full-length product, suggesting that CapRel^SJ46^ was not proteolytically processed (Fig. 3a). Thus, we hypothesized that a specific phage product regulates the C-terminal domain of CapRel^SJ46^ to relieve autoinhibition. To identify such a factor, we sought to identify SECΦ27 mutants that escape CapRel^SJ46^ defense. As no spontaneous escape mutants could be isolated, we used an experimental evolution approach (Fig. 3b). Briefly, we infected cells containing an empty vector or CapRel^SJ46^ with serial dilutions of phage in microtiter plates. After overnight incubation, we collected and pooled the phages from cleared wells, which indicated successful infection, and used these to seed the next round of infections. Initially, cells harboring the empty vector were infected much better, but after 13 rounds, each phage population had evolved to infect both empty vector and CapRel^SJ46^-containing cells similarly (Fig. 3c). We isolated 10 mutant SECΦ27 clones from 5 independently evolved populations and sequenced their genomes. Remarkably, all 10 clones contained a point mutation in the same gene that encodes a hypothetical protein, Gp57, with 9 clones producing the same L114P substitution and one clone yielding an I115F substitution (Fig. 3d).

**Figure 3.**
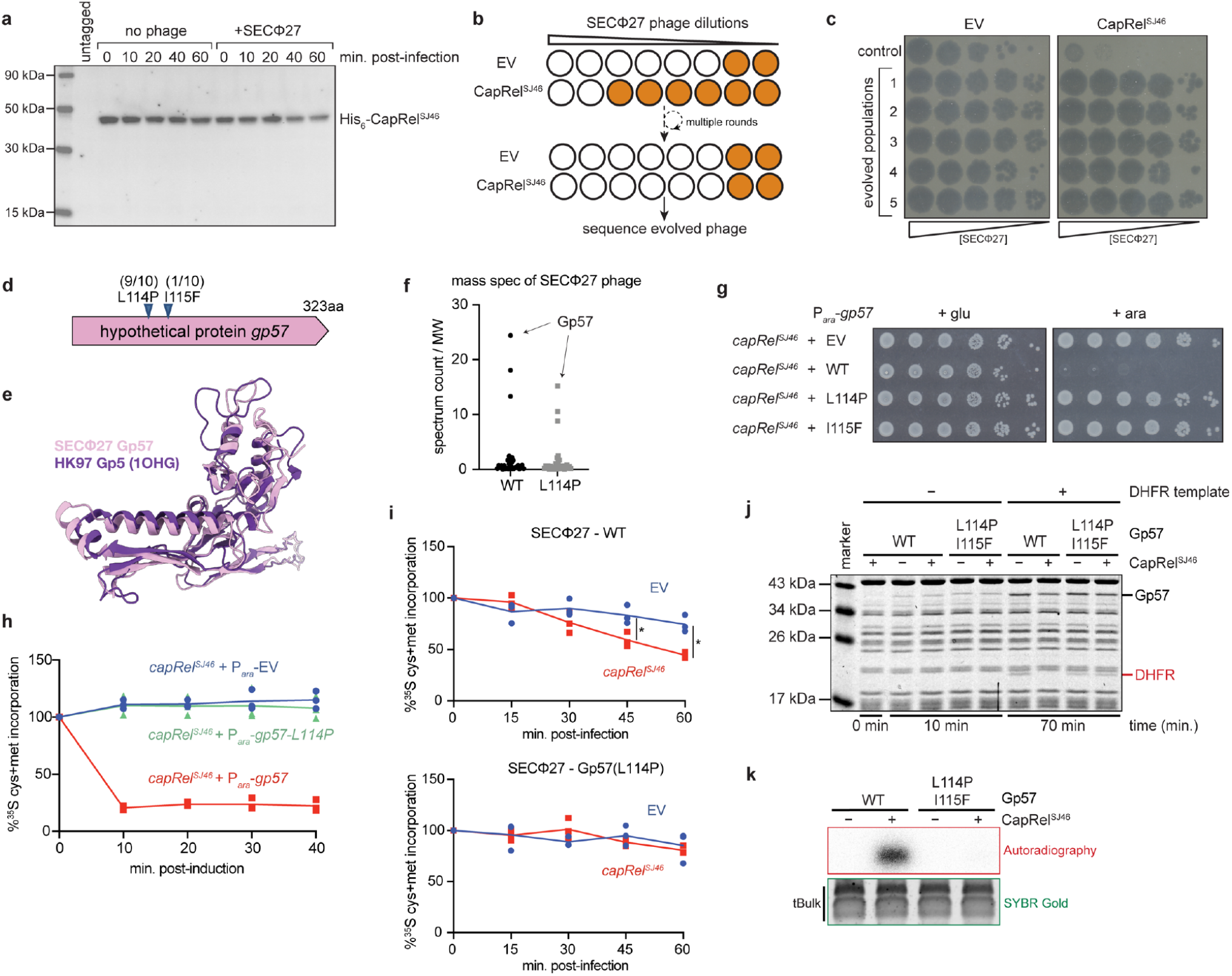
CapRel^SJ46^ is activated by the major capsid protein of SECΦ27 to pyrophosphorylate tRNAs and block translation. (**a**) Immunoblot of His_6_-CapRel^SJ46^ following infection with SECΦ27 (MOI = 100) compared to an uninfected control. (**b**) Schematic of experimental evolution approach used to identify SECΦ27 escape mutants that can infect cells harboring CapRel^SJ46^. (**c**) Serial dilutions of 5 independently evolved populations of SECΦ27 phage and a control population spotted on cells harboring an empty vector (*left*) or CapRel^SJ46^ (*right*). (**d**) Summary of escape mutants identified, all of which map to a hypothetical protein encoded by gene 57 of SECΦ27. (**e**) AlphaFold-predicted structure of Gp57 compared to the major capsid protein Gp5 from phage HK97, which has the eponymous HK97-fold. (**f**) Mass spectrometry analysis of SECΦ27 phage lysates, indicating that the hypothetical protein Gp57 has the highest spectrum count for both WT and an escape mutant producing the L114P variant. (**g**) Serial dilutions on media containing glucose (*left*) or arabinose (*right*) of cells expressing CapRel^SJ46^ from its native promoter and expressing the indicated variant of Gp57 from an arabinose-inducible promoter. (**h**) Cells harboring CapRel^SJ46^ and expressing the wild-type or L114P variant of Gp57 from an arabinose-inducible promoter or harboring an empty vector were pulse-labeled with ^35^S-Cys/Met at the times indicated post-addition of arabinose. (**i**) Same as (**h**) but for cells carrying CapRel^SJ46^ or an empty vector and at times post-infection with SECΦ27 (*top*) or the SECΦ27 escape mutant with the L114P variant of Gp57 (*bottom*) at MOI = 100. Asterisks indicate p < 0.05 (unpaired two-tailed t-test). (**j**) *In vitro* transcription-translation (PURExpress) assays using DHFR production from a DNA template as the readout of expression activity. Purified CapRel^SJ46^ was added to each reaction along with a template for also producing Gp57 (wild-type or the L114P I115F variant). (**k**) Autoradiography of reactions in which purified CapRel^SJ46^ was incubated with [γ-^32^P]-ATP, bulk *E. coli* tRNAs, and Gp57 (WT or the L114P I115F variant). SYBR Gold staining of bulk tRNAs serves as a loading control.

The structure of the hypothetical protein Gp57 predicted by AlphaFold^32^ is highly similar (DALI Z-score of ∼17) to the HK97-fold commonly adopted by major capsid proteins of dsDNA viruses including bacteriophages and Herpesviruses^35^ (Fig. 3e). By performing mass spectrometry on wild-type and escape mutant SECΦ27 phages, we identified this hypothetical protein as the most abundant protein in mature virions, consistent with it being the major capsid protein of SECΦ27 (Fig. 3f and Table S2).

Our results suggested that wild-type Gp57 from SECΦ27 activates CapRel^SJ46^, with the escape mutants preventing activation while retaining the ability to form a capsid. To test this hypothesis, we first examined whether Gp57 alone is sufficient to activate CapRel^SJ46^. Indeed, co-producing wild-type Gp57 with wild-type CapRel^SJ46^ was highly toxic to cells in the absence of phage infection, whereas neither evolved variant (L114P or I115F) of Gp57 had a measurable effect on growth when co-produced with CapRel^SJ46^ (Fig. 3g). As controls, we confirmed that expressing the wild-type or either Gp57 variant was not toxic on its own or if co-produced with a catalytically compromised CapRel^SJ46^ (Fig. S3b).

To examine the basis of CapRel^SJ46^ toxicity we first co-produced it with wild-type or the L114P variant of Gp57 and then measured the effects on bulk transcription and translation by pulse-labeling with ^3^H-uridine and ^35^S-methionine/^35^S-cysteine, respectively. Active CapRel^SJ46^ produced with wild-type Gp57 robustly inhibited translation but not transcription (Fig. 3h and S3c), whereas no effect was seen with Gp57(L114P). Similar effects were seen when overexpressing just the N-terminal domain of CapRel^SJ46^ (Fig. S3d). We also measured bulk translation and transcription following SECΦ27 infection of CapRel^SJ46^-containing cells and observed a decrease in translation but not transcription with wild-type SECΦ27. No effect on translation was seen with the evolved mutant phage producing Gp57(L114P) (Fig. 3i and S3e).

Next, we measured the ability of full-length CapRel^SJ46^ to affect translation *in vitro* using the reconstituted *in vitro* transcription-translation system. Purified CapRel^SJ46^ inhibited synthesis of a control DHFR protein in the presence of the SECΦ27 major capsid protein Gp57, whereas no inhibition was seen for the L114P I115F variant of Gp57 (Fig. 3j). We also incubated wild-type Gp57 or the L114P I115F variant with [γ-^32^P]-ATP and bulk *E. coli* tRNAs in the presence and absence of purified CapRel^SJ46^. Wild-type Gp57 strongly stimulated the pyrophosphorylation of tRNAs by CapRel^SJ46^, like the previously characterized toxSAS enzymes FaRel2 and PhRel2^28^ (Fig. 3k). With the L114P I115F variant of Gp57, tRNA pyrophosphorylation was reduced to the background levels seen with CapRel^SJ46^ alone. Together, our results demonstrate that Gp57, the major capsid protein of SECΦ27, is both necessary and sufficient to activate CapRel^SJ46^, enabling it to pyrophosphorylate tRNAs and inhibit translation.

### CapRel^SJ46^ directly binds to the major capsid protein of SECΦ27

To test whether the SECΦ27 major capsid protein directly binds CapRel^SJ46^, we first immunoprecipitated CapRel^SJ46^-FLAG from cells infected with wild-type phage or the mutant that produces Gp57(L114P) after verifying the tag does not affect CapRel^SJ46^ function (Fig. S3a). We detected Gp57 that had co-precipitated with CapRel^SJ46^ by mass spectrometry when cells were infected with wild-type phage, with a significant reduction in the mutant phage (Fig. S4a-b). In addition, we co-produced CapRel^SJ46^-FLAG and Gp57-HA and found that wild-type, but not the L114P or I115F variant of the capsid protein, co-precipitated with CapRel^SJ46^-FLAG (Fig. 4a and S4c). Finally, we purified both full-length CapRel^SJ46^ and Gp57, and used isothermal titration calorimetry to show that they interact directly with an affinity of 190 nM (Fig. 4b).

**Figure 4.**
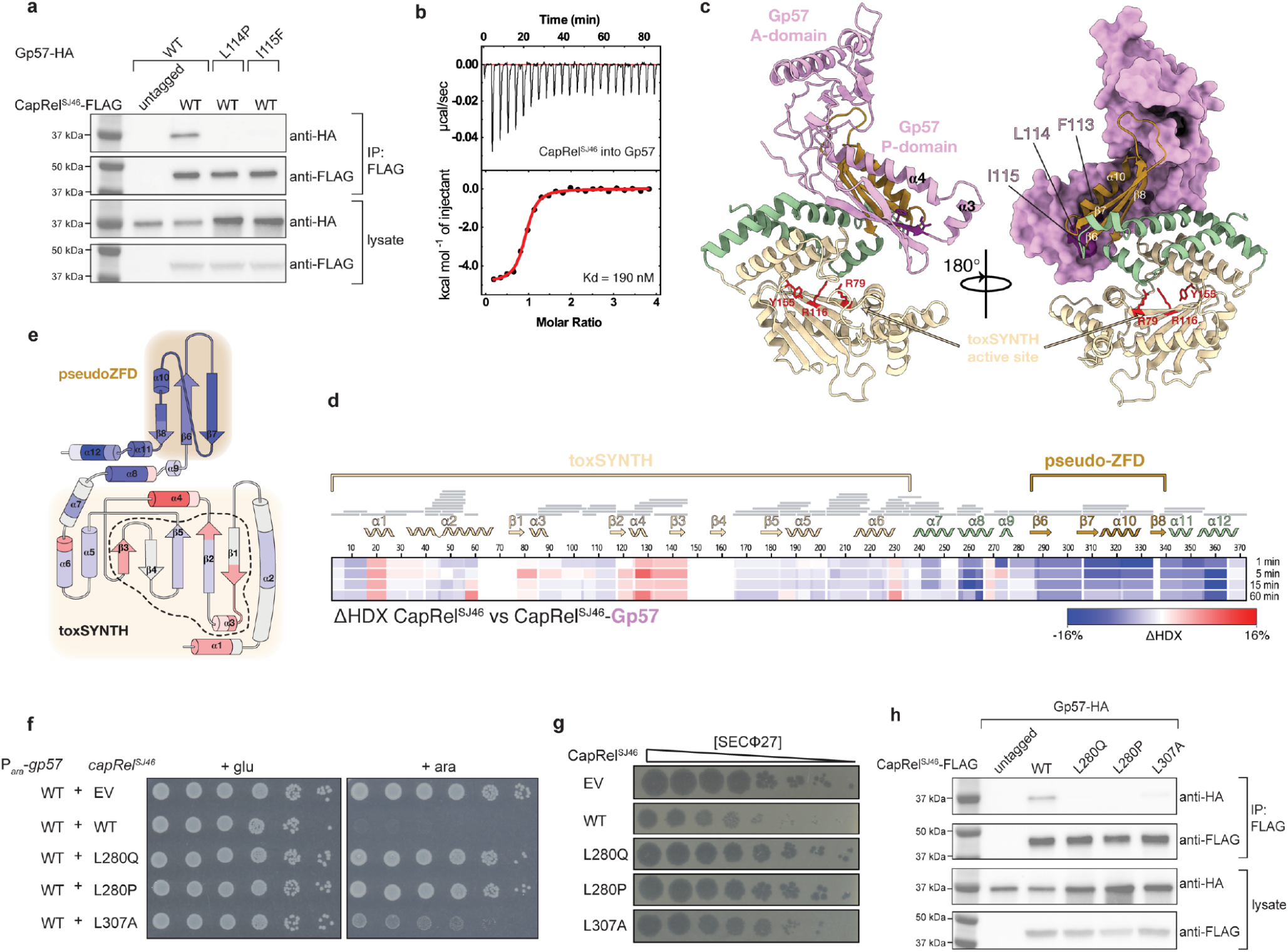
The SECΦ27 major capsid protein Gp57 binds directly to the pseudo-ZFD of CapRel^SJ46^. (**a**) From cells expressing CapRel^SJ46^-FLAG and Gp57-HA (wild-type or mutant variant), the CapRel^SJ46^-FLAG was immunoprecipitated and probed for the presence of the indicated variant of Gp57-HA. Lysates used as input for the IP were probed as controls for expression levels. (**b**) Binding of CapRel^SJ46^ to Gp57 monitored by isothermal titration calorimetry (ITC). (**c**) Structural model of the CapRel^SJ46^-Gp57 complex predicted by AlphaFold. According to the model, the P-domain of Gp57 (in pink) recognizes the pseudo-ZFD (in orange) and anchor regions (in green) of CapRel^SJ46^. This interaction prevents the recoil of pseudo-ZFD to the active site and activates the enzyme. (**d**) “HDX between CapRel^SJ46^ and CapRel^SJ46^-Gp57 displayed as a difference heat map. Red indicates elevated deuteration of CapRel^SJ46^ in the presence of Gp57; blue signi?es lower deuteration. (**e**) Topological representation of CapRel^SJ46^ colored according to the “HDX. The active site of the enzyme is marked by a black dashed outline and the catalytic toxSYNTH domain and the phage-recognition pseudo-ZFD are shadowed in light yellow and light orange. (**f**) Serial dilutions on media containing glucose (*left*) or arabinose (*right*) of cells expressing the indicated mutant of CapRel^SJ46^ from its native promoter and the wild-type Gp57 from an arabinose-inducible promoter. (**g**) Serial dilutions of SECΦ27 phage spotted on cells expressing the indicated mutant of CapRel^SJ46^ or an empty vector. (**h**) Same as in (**a**) but with the indicated mutants of FLAG-CapRel^SJ46^.

Consistent with this tight-binding interaction, the *ab initio* AlphaFold prediction of the CapRel^SJ46^-Gp57 complex has a large contact interface of around 1800 Å^2^ (Fig. 4c). In the complex, CapRel^SJ46^ adopts the same open state seen in our crystal structure (Fig. 2d), with the pseudo-ZFD making extensive contacts with the β-sheet and spine α-helix of the peripheral (P)-domain of Gp57 (Fig. 4c and S4d). Notably, this region of Gp57 contains the residues L114 and I115 identified in our escape mutants. The complex predicted further interactions of pseudo-ZFD β6-β7 loop with the β6-α5 and β8-β9 loops of the axial (A)-domain of Gp57. In this arrangement Gp57 prevents the recoil of pseudo-ZFD to block the active site of the enzyme while stabilizing the YxxY motif in the non-neutralizing hinge conformation.

Hydrogen-deuterium exchange (HDX) monitored by mass spectrometry strongly supported the AlphaFold predictions. In the presence of Gp57, the pseudo-ZFD of CapRel^SJ46^ became more protected with the strongest protection mapping to α10, β8, and the C-terminal α-helical extension (Fig. 4d-e and S4f-g). This overlaps the same region critical for phage specificity (Fig. 2b). The HDX data also confirmed the interface formed between Gp57 P-domain β5 and CapRel^SJ46^ pseudo-ZFD as well as the Gp57 A-domain β8-β9 loop and CapRel^SJ46^ β6-β7 loop. Finally, we observed increased deuterium uptake in CapRel^SJ46^ in residues 110-124 of β2 and 125-130 of α4 which are part of the adenine coordination pocket of toxSYNTH, thus confirming that interaction with Gp57 exposes the active site of the enzyme (Fig. 4d-e and S4f-g).

To further validate the role of the pseudo-ZFD in binding and activating CapRel^SJ46^, we performed error-prone PCR-based mutagenesis on this domain and screened for mutations that disrupted activation of CapRel^SJ46^ when it was co-produced with the capsid protein Gp57. The substitutions L280Q and L280P drastically reduced the toxicity of CapRel^SJ46^ in the presence of wild-type Gp57 (Fig. 4f), and prevented CapRel^SJ46^ from protecting against SECΦ27 infection (Fig. 4g). Importantly, these CapRel^SJ46^ variants still protected *E. coli* against phage T2 and T4, indicating that these variants retained structural integrity (Fig. S4e). The substitution L307A had similar, but reduced, effects on CapRel^SJ46^ activity (Fig. 4f-g).

The crystal structure of CapRel^SJ46^ suggested that L280 and L307 in the wild-type protein promote the open, active state, with L280 stabilizing one of the hinge regions involving the pseudo-ZFD and L307 structuring the β6-β7 loop that interacts with Gp57 A-domain. The L280Q and L280P variants of CapRel^SJ46^ were unable to co-precipitate the major capsid protein of SECΦ27, and the L307A substitution significantly reduced binding in this assay (Fig. 4h). In sum, our findings strongly support a model in which the C-terminal pseudo-ZFD of CapRel^SJ46^ directly recognizes the major capsid protein of SECΦ27, thereby triggering a relief of autoinhibition of the N-terminal toxSYNTH domain.

### CapRel^SJ46^ can be activated by capsid homologs of other phages

The pseudo-ZFD of CapRel^SJ46^, including residues L280 and L307, is the least well conserved portion of the protein (Fig. S2a and S2f). This variability may reflect a Red Queen dynamic, a hallmark of many host-pathogen interfaces that arises from cycles of selective pressure on pathogens to evade host immunity followed by selection on host immune factors to restore recognition of a pathogen^5^. As triggers of the CapRel defense system, phage capsid proteins are likely under pressure to diversify, while retaining the ability to form a capsid, leading to a selective pressure on the pseudo-ZFD of CapRel to diversify and retain its interaction with the capsid proteins. To test this hypothesis, we examined three phages from the BASEL collection^36^ (Bas4, Bas5 and Bas8) that are closely related to SECΦ27 and contain a close homolog of Gp57 called Gp8 (Fig. S5a). We first found that co-expressing the major capsid homologs from Bas5 and Bas8, but not that of Bas4, with CapRel^SJ46^ rendered CapRel^SJ46^ toxic, as with the SECΦ27 capsid protein. We then tested whether CapRel^SJ46^ protects against these phages and found that it protected strongly against Bas5 and Bas8, but not Bas4 (Fig. 5b).

**Figure 5.**
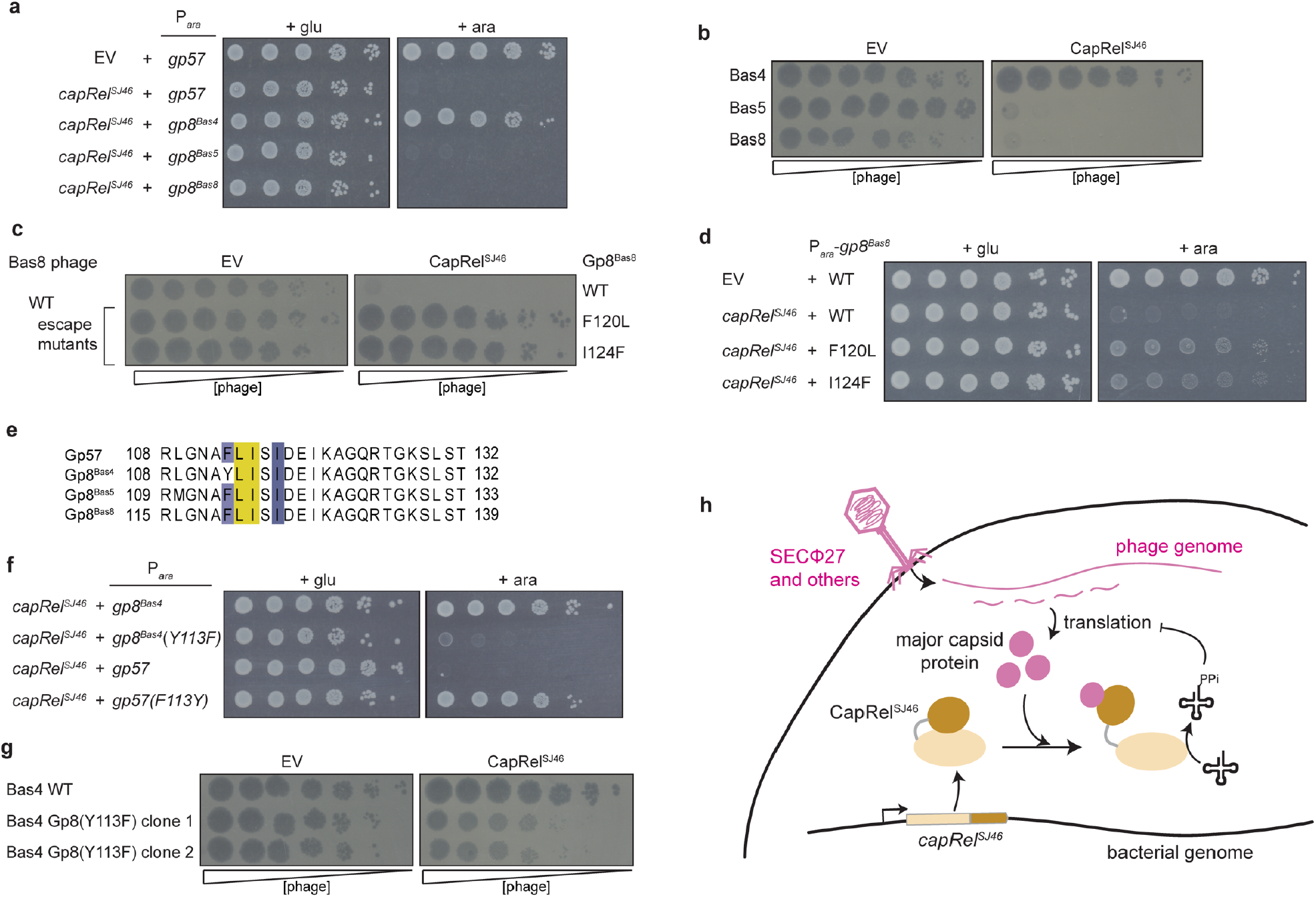
Evidence for the coevolution of CapRel^SJ46^ and the major capsid protein of SECΦ27 and related phages. (**a**) Serial dilutions on media containing glucose (*left*) or arabinose (*right*) of cells expressing CapRel^SJ46^ from its native promoter and the major capsid protein homolog from the phage indicated via an arabinose-inducible promoter. (**b**) Serial dilutions of the phages indicated spotted on lawns of cells harboring CapRel^SJ46^ or an empty vector. (**c**) Serial dilutions of wild-type Bas8 phage or the escape mutants bearing the major capsid mutations indicated spotted on lawns of cells harboring CapRel^SJ46^ or an empty vector. (**d**) Serial dilutions on media containing glucose (*left*) or arabinose (*right*) of cells expressing CapRel^SJ46^ from its native promoter or harboring an empty vector and producing the indicated variant of the Bas8 major capsid protein from an arabinose-inducible promoter. (**e**) Alignment of the region of the major capsid protein in SECΦ27, Bas5, and Bas8 that triggers CapRel^SJ46^, along with Bas4 which has a tyrosine at position 113 instead of phenylalanine. (**f**) Serial dilutions on media containing glucose (*left*) or arabinose (*right*) of cells expressing CapRel^SJ46^ from its native promoter and the Bas4 or SECΦ27 major capsid protein variant indicated from an arabinose-inducible promoter. **(g)** Serial dilutions of wild-type Bas4 or two mutant clones containing Y113F in the major capsid protein Gp8 spotted on lawns of cells harboring CapRel^SJ46^ or an empty vector. (**h**) Model for the direct activation of CapRel^SJ46^ by the major capsid protein of SECΦ27 and related phages. After genome injection, the production of the major capsid protein triggers relief of autoinhibition by the C-terminal antitoxin of CapRel^SJ46^, leading to pyrophosphorylation of tRNAs by activated CapRel^SJ46^, which inhibits translation and restricts viral infection.

To validate that defense against Bas8 requires activation of CapRel^SJ46^ by this phage’s capsid protein homolog, we isolated spontaneous mutants of Bas8 that escaped defense. Two mutant clones of Bas8 were no longer defended against by CapRel^SJ46^ and contained either an F120L or I124F substitution in the major capsid homolog (Fig. 5c). Both substitutions significantly reduced the capsid protein’s ability to activate CapRel^SJ46^ when co-produced (Fig. 5d). Notably, these two positions were close to the positions of the escape mutants identified in SECΦ27 Gp57, further confirming that this region in the major capsid protein is important for activating CapRel^SJ46^ (Fig. 5e).

Unlike Bas8, Bas4 was not defended against by CapRel^SJ46^ and its capsid homolog did not activate CapRel^SJ46^ despite being 98% identical to SECΦ27 Gp57, with just 5 amino acid differences between the two. However, one difference is at position 113, near the region that likely binds to CapRel^SJ46^. This residue is a phenylalanine in SECΦ27, Bas5, and Bas8, but a tyrosine in Bas4 (Fig. 5e). We tested whether this residue is critical for activation by making a Y113F substitution in the Bas4 capsid homolog and found that it gained the ability to activate CapRel^SJ46^ when coproduced (Fig. 5f). Conversely, a F113Y substitution in the SECΦ27 capsid protein abolished its ability to activate CapRel^SJ46^. Additionally, we mutated Bas4 phage such that it produces major capsid protein harboring the Y113F substitution. This mutant phage could still produce mature virions, but was now defended against by CapRel^SJ46^ (Fig. 5g). These results support the notion of a Red Queen dynamic between the pseudo-ZFD of CapRel and the phage capsid proteins that directly bind and activate CapRel.

## Conclusions

We propose the following model for CapRel^SJ46^ activation by SECΦ27 (Fig. 5h). Without phage infection, CapRel^SJ46^ adopts an inactive, closed conformation in cells with its C-terminal antitoxin domain autoinhibiting the N-terminal toxin domain. Upon infection, the major phage capsid protein is produced and directly binds to CapRel^SJ46^ to stabilize the active, open state. This open state enables CapRel^SJ46^ to pyrophosphorylate tRNAs and inhibit translation, leading to an abortive infection that prevents propagation of phage through a population of cells. Importantly, our results imply that type II TA systems, which feature protein antitoxins, can be activated without proteolysis of the antitoxin, which is often asserted as their primary means of activation.

Major capsid proteins, like Gp57 from SECΦ27, may be a common trigger for both TA systems and other anti-phage defense systems. Prior studies found that a short peptide called Gol within the major capsid protein Gp23 of T4 can activate the Lit protease in *E. coli* if both components are overproduced^37,38^. For PifA, which allows the F plasmid to exclude T7, escape mutants mapped to the major capsid protein, but this interaction has not been studied biochemically^39^. Recent work reported that mutations in the major capsid protein of T5 allow it to overcome Pycsar-mediated defense, but the capsid protein alone is insufficient to activate Pycsar^40^. We anticipate that major capsid proteins may emerge as common, direct triggers for a diverse range of anti-phage defense systems. As with PAMPs in eukaryotes, relying on an essential, abundant component of phages for activation may help ensure that an immune response is only mounted following an infection. Notably, the capsid proteins of some eukaryotic viruses stimulate mammalian innate immune pathways. For instance, HIV capsid protein is directly detected in the host cell cytoplasm and nucleus by TRIM5 and NONO, respectively, to trigger innate immune activation^7,9^. Thus, our results suggest that similar principles of pathogen detection underlie the function and molecular basis of innate immunity in all domains of life.

## Acknowledgements

We thank A. Harms for generously sharing the BASEL phage collection, the MIT BioMicro Center and its staff for their support in sequencing, the MIT Biopolymers & Proteomics Core and its staff for their help in mass spectrometry experiments. We thank K. Gozzi and B. Wang for comments on the manuscript and all members of the Laub lab for helpful discussions. G.C.A and V.H. were supported by the Swedish Research council (grant 2018-00956 within the RIBOTARGET consortium under the framework of JPIAMR, project grants 2017-03783 and 2021-01146 to V.H., project grant 2019-01085 to G.C.A.), the Knut and Alice Wallenberg Foundation (2020.0037 to G.C.A), the Ragnar Söderberg Foundation (M23/14 to V.H.), the European Regional Development Fund through the Centre of Excellence for Molecular Cell Technology (V.H.), and the Estonian Science Foundation (project grant PRG335 to V.H.). A.G-P. was supported by Fonds National de Recherche Scientifique (FRFS-WELBIO CR-2017S-03, FNRS CDR J.0068.19, FNRS-EQP UN.025.19 and FNRS-PDR T.0090.22), the European Research Council (CoG DiStRes, n° 864311), the Joint Programming Initiative on Antimicrobial Resistance (JPI-EC-AMR-R.8004.18), the Programme Actions de Recherche Concerté 2016-2021, Fonds Jean Brachet and the Fondation Van Buuren, Chargé de Recherches fellowship from the FNRS n° CR/DM-392 (HeT), the European Union’s Horizon 2020 research and innovation programme under the Marie Skłodowska-Curie grant agreement N° 801505 (IF@ULB postdoctoral grant to A.A.). K.C.W. is a fellow of the FRIA, C.M. is supported as a Research Associate of the F.R.S.-F.N.R.S. The authors acknowledge the use of the PROXIMA 1 and 2A beamlines at the Soleil synchrotron (Gif-sur-Yvette, France). M.T.L. is an Investigator of the Howard Hughes Medical Institute.

## Author Contributions

Experiments were conceived and designed by T.Z., T.K., G.A., V.H., A.G-P., M.T.L. Phage and bacterial experiments, as well as incorporation and co-IP assays, were done by T.Z. with assistance from M.L. and S.S. Metabolic labeling experiments were done by T.Z. and T.B. Cell-free translation and tRNA pyrophosphorylation assays were done by T.Z and T.K. CapRel purification was done by T.Z., T.K., H.T., and A.T. ITC was performed by H.T., A.T., and A.C. HDX was performed by C.M. and K.C.W. X-ray data collection and analyses was performed by H.T., A.T., and A.G-P. Bioinformatic analyses were performed by T.Z. and G.C.A. Figure design, manuscript writing, and editing done by T.Z., T.K., G.A., V.H., A.G-P., M.T.L. Project supervision and funding provided by G.A., V.H., A.G-P., M.T.L.

## Author Information

The authors declare no competing financial interests. Correspondence and requests for materials should be addressed to V.H., A.G-P., and M.T.L.

## Data and materials availability

Structural data are available in PDB (7ZTB). HDX data raw data can be accessed at: doi.org/10.6084/m9.figshare.19745089. Sequencing data are available in the Sequence Read Archive (SRA) under BioProject PRJNA837951. All other data are available in the manuscript or the supplementary materials. Materials including strains and plasmids are available upon reasonable request.

## Methods

### Strains and growth conditions

All bacterial and phage strains used in this study are listed in Table S3. *Escherichia coli* strains were routinely grown at 37 °C in Luria broth (LB) medium for cloning and maintenance. Phages were propagated by infecting a culture of *E. coli* MG1655 at an OD_600_ ∼0.1-0.2 with a MOI of 0.1. Cleared cultures were pelleted by centrifugation to remove residual bacteria and filtered through a 0.2 µm filter. Chloroform was then added to phage lysates to prevent bacterial growth. All phage infection experiments in liquid media and phage spotting experiments were performed in LB medium at 25 °C, except for spotting of T2 and T4 on strains producing CapRel^SJ46^ variants, which was performed in M9 medium (6.4 g/L Na_2_HPO_4_-7H_2_O, 1.5 g/L KH_2_PO_4_, 0.25 g/L NaCl, 0.5 g/L NH_4_Cl medium supplemented with 0.1% casamino acids, 0.4% glycerol, 0.4% glucose, 2 mM MgSO_4_, and 0.1 mM CaCl_2_) at 30 °C. For liquid induction experiments from pBAD33 vectors, bacterial cells were grown in M9 medium. Antibiotics were used at the following concentrations (liquid; plates): carbenicillin (50 μg/mL; 100 μg/mL), chloramphenicol (20 μg/mL; 30 μg/mL).

### Plasmid construction

All plasmids are listed in Table S4. All primers and synthesized gene sequences are listed in Table S5.

pBR322-*capRel* constructs: DNA encoding *capRel*^*SJ46*^, *capRel*^*Ebc*^, and *capRel*^*Kp*^ open reading frames were codon-optimized for expression in *E. coli* and 100-200 bp of the upstream region from the source organism was added in each case for native expression (TZ-1 to TZ-5). DNA was commercially synthesized by Integrated DNA Technology as gBlocks and assembled into a promoter-less backbone of pBR322 amplified with TZ-6 and TZ-7 by Gibson assembly. Mutations that produce the single amino-acid substitutions CapRel^SJ46^(Y155A), CapRel^Ebc^(Y153A), CapRel^SJ46^(L280Q), CapRel^SJ46^(L280P) and CapRel^SJ46^(L307A) were generated by site-directed mutagenesis using primers TZ-8 to TZ-11 and TZ49 to TZ-54. To add an N-terminal His_6_-tag or a C-terminal FLAG-tag to CapRel^SJ46^, primers TZ41 and TZ-42 or TZ45 and TZ46 were used to PCR-amplify pBR322-*capRel*^*SJ46*^ followed by Gibson assembly. pBR322-*capRel-chimera* was constructed by inserting *capRel*^*Ebc*^(270-339) that had been PCR*-* amplified with TZ-22 and TZ-23 into pBR322-*capRel*^*SJ46*^ linearized with TZ-20 and TZ-21 using Gibson assembly.

pBAD33-*capRel*^*SJ46*^ constructs: *capRel*^*SJ46*^*(1-272)* or full-length *capRel*^*SJ46*^ was PCR-amplified with TZ14 and TZ15, or TZ14 and TZ24, respectively, and inserted into pBAD33 linearized with TZ-12 and TZ-13 using Gibson assembly. pBAD33-*capRel*^*SJ46*^ variants (A77K, R116A, V338A, L339A, A341K, A351K, Y352A or Y355A) were constructed by site-directed mutagenesis using primers TZ25 to TZ40. pBAD33-*capRel*^*SJ46*^ variants (R78A, K311A, R314A, E319A, K346A) were constructed by site-directed mutagenesis using primers TZ75 to TZ84.

pEXT20-*capRel*^*SJ46*^ construct: *capRel*^*SJ46*^*(273-373)* was PCR-amplified with primers TZ-18 and TZ-19, and then inserted into linearized pEXT20 with TZ-16 and TZ-17 using Gibson assembly.

pBAD33*-gp57* constructs: wild-type or mutant variant (L114P or I115F) *gp57* was PCR-amplified from the corresponding wild-type or escape mutant SECΦ27 phage using primers TZ-43 and TZ-44, and inserted into linearized pBAD33 using Gibson assembly. A C-terminal HA-tag was added to wild-type or mutant *gp57* using primers TZ-47 and TZ-48 to PCR-amplify the corresponding construct followed by Gibson assembly. The F113Y variant of *gp57* was generated by site-directed mutagenesis using primers TZ-63 and TZ-64.

pBAD33-*gp8*: the genes encoding the major capsid protein homologs Gp8^Bas4^, Gp8^Bas5^, and Gp8^Bas8^ were PCR-amplified from the corresponding phage using primers TZ-55 to TZ-60 and inserted into linearized pBAD33 by Gibson assembly. The Y113F variant of *gp8*^*Bas4*^ was generated by site-directed mutagenesis using primers TZ-61 and TZ-62. The F120L and I124F variants of *gp8*^*Bas8*^ were cloned from the corresponding phage escape mutants using primers TZ-59 and TZ-60.

pET-*gp57* constructs: *gp57* and *gp57(L114P I115F)* fragments were PCR-amplified with primers TZ-65 and TZ-66 and either TZ-67 template (for *gp57*) or TZ-68 (for *gp57(L114P I115F)*).

Using Gibson assembly, the resultant linear DNA fragments were inserted into linearized pET24d (without tag) using TZ-69 and TZ-70. Templates TZ-67 and TZ-68 we synthesized as gBlocks by Integrated DNA Technology.

pET24d-*His*_*10*_*-SUMO-capRel*^*SJ46*^ constructs: *capRel*^*SJ46*^ ORF was PCR-amplified using primers TZ-71 and TZ-72 as well as pBAD-*capRel*^*SJ46*^ as template, and, using Gibson assembly, inserted into a linearized pET24d-*His*_*10*_-*SUMO* plasmid using primers TZ-73 and TZ-74.

### Strain construction

Plasmids described above were introduced into *E. coli* MG1655 or BW27783 by TSS transformation or electroporation.

Bas4 mutant phage were generated using a CRISPR-Cas system for targeted mutagenesis as described previously^41^. Briefly, sequences for RNA guides to target Cas9-mediated cleavage were designed using the toolbox in Geneious Prime 2021.2.2 and selected for targeting of *gp8*^*Bas4*^ but nowhere else in the Bas4 genome. The guides were inserted into the pCas9 plasmid and tested for their ability to restrict Bas4. An efficient guide was selected and the pCas9-guide plasmid was co-transformed into *E. coli* MG1655 with a high copy-number repair plasmid containing *gp8*^*Bas4*^*(Y113F)* with the guide mutated to prevent self-cutting. The wild-type Bas4 phage was plated onto a strain containing both the pCas9-*guide* and the repair plasmid, and single plaques were screened by Sanger Sequencing. Two clones that produce the Y113F substituted Gp8 were propagated twice on strains containing only pCas9-*guide* for further selection and genomes were sequence verified by Illumina sequencing as described below.

### Toxicity assays on solid media

For producing the CapRel^SJ46^ N- and C-terminal domains, single colonies of *E. coli* MG1655 containing pBAD33-*capRel*^*SJ46*^*(1-272)* and pEXT20-*capRel*^*SJ46*^*(273-373)* or the corresponding empty vectors were grown for 6 hours at 37 °C in LB-glucose to saturation. 200 µL of each saturated culture was then pelleted by centrifugation at 4000 *g* for 10 min, washed once in 1x phosphate-buffered saline (PBS), and resuspended in 400 µL 1x PBS. Cultures were then serially-diluted 10-fold in 1x PBS and spotted on M9L plates (M9 medium supplemented with 5% LB (v/v)) further supplemented with 0.4% glucose, 0.2% arabinose or 0.2% arabinose and 100 µM IPTG. Plates were then incubated at 37 ºC overnight before imaging.

For producing full-length CapRel^SJ46^, *E. coli* MG1655 containing pBAD33-*capRel*^*SJ46*^ or a mutant form of *capRel*^*SJ46*^ were grown to saturation and processed as above. Cultures were plated onto 0.4% glucose and 0.2% arabinose and incubated at 37 ºC overnight.

For co-producing CapRel^SJ46^ and the major capsid proteins from SECΦ27, Bas4, Bas5, or Bas8, *E. coli* MG1655 harboring pBR322-*capRel*^*SJ46*^ and pBAD33-*capsid protein* were grown to saturation and processed as above. Cultures were plated onto 0.4% glucose and 0.2% arabinose and incubated at 37 ºC overnight.

For co-producing CapRel^SJ46^ and variants of the major capsid protein from Bas8, *E. coli* BW27783 harboring pBR322-*capRel*^*SJ46*^ and pBAD33-*gp8*^*Bas8*^ (wild-type or a mutant variant) were grown to saturation and processed as above. Cultures were plated onto 0.4% glucose and 0.0002% arabinose and incubated at 37 ºC overnight.

### Phage spotting assays and efficiency of plaquing (EOP) measurements

Phage stocks isolated from single plaques were propagated in *E. coli* MG1655 at 37 ºC in LB. To titer phage, dilutions of stocks were mixed with *E. coli* MG1655 and melted LB + 0.5% agar and spread on LB + 1.2% agar plates and incubated at 37 ºC overnight. For phage spotting assays, 40 µL of a bacterial strain of interest was mixed with 4 mL LB + 0.5% agar and spread on an LB + 1.2% agar + antibiotic plate. Phage stocks were then serially diluted in 1x FM buffer (20 mM Tris-HCl pH 7.4, 100 mM NaCl, 10 mM MgSO_4_), and 2 µL of each dilution was spotted on the bacterial lawn. Plates were then incubated at 25 ºC overnight before imaging. Efficiency of plaquing (EOP) was calculated by comparing the ability of the phage to form plaques on an experimental strain relative to the control strain. Experiments were replicated 3 times independently and representative images are shown.

For spotting phage T2 and T4 on strains producing CapRel^SJ46^ variants, 40 µL of a bacterial strain of interest was mixed with 4 mL M9 + 0.5% agar and spread on an M9 + 1.2% agar + antibiotic plate. Phage were serially diluted and spotted as described above. Plates were then incubated at 30 ºC overnight before imaging.

### Growth curves following phage infection in liquid culture

Single colonies of *E. coli* MG1655 pBR322-EV or pBR322-*capRel*^*SJ46*^ or pBR322-*capRel*^*SJ46*^*(Y155A)* were grown in LB overnight. Cultures were then back-diluted to OD_600_ = 0.1 in fresh LB and 100 µL cells were added into each well of a 96-well plate. 10 µL of serial-diluted T4 phage were added to each well at the indicated MOI and growth following phage infection was measured at 15 min intervals with orbital shaking at 25 ºC on a plate reader (Biotek). Data reported are the mean and standard deviation of 8 plate replicates and the growth curve experiment was replicated 3 times independently.

### One-step growth curves

Single colonies of *E. coli* MG1655 pBR322-EV or pBR322-*capRel*^*SJ46*^ were grown overnight in LB. Overnight cultures were back-diluted to OD_600_ = 0.05 in 25 mL fresh LB and grown to OD_600_ ∼ 0.3 at 25 ºC. 10 mL of each culture were infected with T4 phage at an MOI of 0.05 in LB at 25 ºC and phages were allowed to adsorb for 10 min before serial dilution in LB three times (1:100, 1:10, 1:10 serial dilution) to three flasks. Then, at indicated time points, 100 µL of infected cells from the corresponding dilution flask were mixed with 100 µL of indicator cells MG1655 pBR322-EV (OD_600_ ∼ 0.3), and the mixtures were mixed with 4 mL of LB + 0.5% agar and spread on LB + 1.2% agar plates. Plates were incubated overnight at 25 ºC and plaques were enumerated the following day. Plaque forming units (pfu/mL) were calculated based on the dilution flask samples were taken from. Data reported are the mean and individual data points from 3 biological replicates.

### Western blot of CapRel^SJ46^ after phage infection

Single colonies of *E. coli* MG1655 pBR322-*His*_*6*_*-capRel*^*SJ46*^ were grown overnight in LB. Overnight cultures were back-diluted to OD_600_ = 0.05 in 25 mL fresh LB and grown to OD_600_ = 0.2 at 25 ºC. Cells were infected with phage SECΦ27 at MOI = 100, and incubated at 25 ºC during the experiment. At each indicated time point (0, 10, 20, 40, 60 min), OD_600_ was measured and 1 mL of cells was pelleted at 21,000 *g* for 2 min at 4 ºC. Supernatant was removed and pellets were flash-frozen in liquid nitrogen. Pellets were thawed and resuspended in 1x Laemmli sample buffer (Bio-Rad) supplemented with 2-mercaptoethanol with OD_600_ normalized. Samples were then boiled at 95 ºC and analyzed by 12% SDS-PAGE and transferred to a 0.45 µm PVDF membrane. Anti-His_6_ antibody (Invitrogen) was used at a final concentration of 1:1000, and SuperSignal West Femto Maximum Sensitivity Substrate (ThermoFisher) was used to develop the blots. Blots were imaged by a ChemiDoc Imaging system (Bio-Rad). Image shown is a representative of 2 independent biological replicates.

### Isolation of phage escape mutants to infect CapRel^SJ46^

The phage evolution experiment was conducted as described previously^42^. Briefly, five independent populations were evolved in a 96-well plate containing a sensitive host *E. coli* MG1655 pBR322-EV and a resistant host *E. coli* MG1655 pBR322-*capRel*^*SJ46*^. One control population was evolved with only the sensitive host. Overnight bacterial cultures were back-diluted to OD_600_ = 0.1 in LB and 100 µL were seeded into each well. Cells were infected with 10-fold serial dilutions of SECΦ27 phage with MOI from 100 to 10^−4^, with one well uninfected to monitor for contamination. Plates were sealed with breathable plate seals and incubated at 25 °C for 6 hours in a plate shaker at 1000 rpm. Cleared wells from each population were pooled, pelleted at 4000 *g* for 20 min to remove bacteria, and the supernatant lysates were transferred to a 96 deep-well block with 40 µL chloroform added to prevent bacterial growth. Lysates were spotted onto both sensitive and resistant hosts to check the defense phenotype. Thirteen rounds of evolution were performed to allow all five populations to overcome CapRel^SJ46^ defense. Evolved clones from each evolved population were isolated by plating to single plaques on lawns of resistant host, and control clones from the control population were isolated on a lawn of the sensitive host. Two clones from each population were propagated using the corresponding host and sequenced as described below.

Bas8 escape mutants were isolated by plating a population of phage onto CapRel^SJ46^-containing cells. 20 µL of 10^11^ pfu/mL Bas8 phage mixed with 40 µL overnight culture of *E. coli* MG1655 pBR322-*capRel*^*SJ46*^ were added to 4 mL of LB + 0.5% agar and spread onto LB + 1.2% agar. Plates were incubated at 25 °C overnight. Single plaques were isolated and propagated using the same strain in LB at 25 °C. Amplified phage lysates were pelleted to remove bacteria, and then plated to single plaques and propagated similarly for a second round of isolation to improve purity and sequenced as described below.

### Phage DNA extraction and Illumina sequencing

To extract phage DNA, high titer phage lysates (> 10^6^ pfu/µL) were treated with DNase I (0.001 U/µL) and RNase A (0.05 mg/mL) at 37 °C for 30 min. 10 mM EDTA was used to inactivate the nucleases. Lysates were then incubated with Proteinase K at 50 °C for 30 min to disrupt capsids and release phage DNA. Phage DNA was isolated by ethanol precipitation. Briefly, NaOAc pH 5.2 was added to 300 mM followed by 100% ethanol to a final volume fraction of 70%. Samples were incubated at −80 °C overnight, pelleted at 21,000 *g* for 20 min and supernatant removed. Pellets were washed with 100 µL isopropanol and 200 µL 70% (v/v) ethanol, and then aired dried at room temperature and resuspended in 25 µL 1x TE buffer (10 mM Tris-HCl, 0.1 mM EDTA, pH = 8). Concentrations of extracted DNA were measured by NanoDrop (Thermo Fisher Scientific).

To prepare Illumina sequencing libraries, 100-200 ng of genomic DNA was sheared in a Diagenode Bioruptor 300 sonicator water bath for 20x 30 s cycles at maximum intensity. Sheared genomic DNA was purified using AmpureXP beads, followed by end repair, 3’ adenylation, and adaptor ligation. Barcodes were added to both 5’ and 3’ ends by PCR with primers that anneal to the Illumina adaptors. The libraries were cleaned by Ampure XP beads using a double cut to elute fragment sizes matching the read-lengths of the sequencing run. Libraries were sequenced on an Illumina MiSeq at the MIT BioMicro Center. Illumina reads were assembled to the reference genomes using Geneious Prime 2021.2.2.

### Mass spectrometry of phages

Wild-type or mutant (L114P in Gp57, evolved clone 1 from population 3) SECΦ27 phage were propagated in *E. coli* MG1655 for high titer stocks. Briefly, *E. coli* MG1655 (OD_600_ = 0.2) in LB were infected with phages at MOI = 0.1 and incubated at 37 °C for 4 hours. Cells were pelleted at 4000 *g* for 10 min and supernatant lysates were filtered through 0.2 µm filters. 500 µL of phage stocks (10^10^ pfu/µL) were further concentrated with Amicon Ultra filter (MW 100 kDa) and washed twice with 1x FM buffer (20 mM Tris-HCl pH 7.4, 100 mM NaCl, 10 mM MgSO_4_). Concentrated phage lysates were boiled to denature virions and run on 4-20% SDS-PAGE. Each lane from the gel was excised. Proteins were reduced with 10 mM dithiothreitol (Sigma) for 1 hour at 56 °C and then alkylated with 20 mM iodoacetamide (Sigma) for 1 hour at 25 °C in the dark. Proteins were then digested with 12.5 ng/µL modified trypsin (Promega) in 50 µL 100 mM ammonium bicarbonate, pH 8.9 at 25 °C overnight. Peptides were extracted by incubating the gel pieces with 50% acetonitrile/5% formic acid then 100 mM ammonium bicarbonate, repeated twice followed by incubating the gel pieces with 100% acetonitrile then 100 mM ammonium bicarbonate, repeated twice. Each fraction was collected, combined, and reduced to near dryness in a vacuum centrifuge. Peptides were desalted using Pierce Peptide Desalting Spin Columns (Thermo) and then lyophilized. The tryptic peptides were separated by reverse phase HPLC (Thermo Ultimate 3000) using a Thermo PepMap RSLC C18 column over a 90 min gradient before nano-electrospray using an Exploris mass spectrometer (Thermo). Solvent A was 0.1% formic acid in water and solvent B was 0.1% formic acid in acetonitrile. Detected peptides were mapped to SECΦ27 protein sequences and the abundance of proteins were estimated by number of spectrum counts/molecular weight (SC/MW) to normalize for protein sizes.

### Co-immunoprecipitation (co-IP) analysis

For immunoprecipitation of CapRel^SJ46^ after phage infection, *E. coli* MG1655 containing pBR322-*capRel*^*SJ46*^*-FLAG* were grown overnight in LB. Overnight cultures were back-diluted to OD_600_ = 0.05 in 175 mL of LB and grown to OD_600_ ∼ 0.3 at 25 ºC. Cells were infected with wild-type or mutant (L114P in Gp57, evolved clone 1 from population 3) SECΦ27 at MOI = 100 and incubated at 25 ºC. At the indicated time points (15 min or 40 min), OD_600_ was measured and 50 mL of cells were pelleted at 6000 *g* for 5 min at 4 ºC. Uninfected cells were harvested at 0 min before phage infection. Supernatant was removed and cells were resuspended in 900 μL lysis buffer (25 mM Tris-HCl, 150 mM NaCl, 1 mM EDTA, 1% Triton X-100 and 5% glycerol) supplemented with protease inhibitor (Roche), 1 μL/mL Ready-Lyse™ Lysozyme Solution (Lucigen) and 1 μL/mL benzonase nuclease (Sigma). Samples were lysed by two freeze-thaw cycles, and lysates were normalized by OD_600_. Lysates were pelleted at 21,000 *g* for 10 min at 4 °C, and 850 μL of supernatant were incubated with pre-washed anti-FLAG M2 magnetic beads (Sigma) beads for 1 hour at 4 ºC with end-over-end rotation. Beads were then washed 3 times with lysis buffer containing 350 mM NaCl but free of detergent. On-bead reduction, alkylation and digestion were performed. Proteins were reduced with 10 mM dithiothreitol (Sigma) for 1 hour at 56 °C and then alkylated with 20 mM iodoacetamide (Sigma) for 1 hour at 25 °C in the dark. Proteins were then digested with modified trypsin (Promega) at an enzyme/substrate ratio of 1:50 in 100 mM ammonium bicarbonate, pH 8 at 25 °C overnight. Trypsin activity was halted by addition of formic acid (99.9 %, Sigma) to a final concentration of 5 %. Peptides were desalted using Pierce Peptide Desalting Spin Columns (Thermo) then lyophilized. The tryptic peptides were subjected to LC-MS/MS as described above. Experiments were performed 2 times independently and spectral counts are reported. Ratio of spectral counts between Gp57 and CapRel^SJ46^ were calculated and graphed for normalization.

For co-producing CapRel^SJ46^ and Gp57, *E. coli* MG1655 containing pBR322-*capRel*^*SJ46*^ or pBR322-*capRel*^*SJ46*^*-FLAG* (wild type or mutants) and pBAD33-*gp57-HA* (wild type or mutants) were grown overnight in M9-glucose. Overnight cultures were back-diluted to OD_600_ = 0.05 in 50 mL of M9 (no glucose) and grown to OD_600_ ∼ 0.3 at 37 ºC. Cells were induced with 0.2% arabinose for 30 min at 37 ºC, then OD_600_ was measured and cells were pelleted at 4000 *g* for 10 min at 4 ºC. Supernatant was removed and cells were resuspended in 900 μL lysis buffer as described above. Samples were lysed by two freeze-thaw cycles, and lysates were normalized by OD_600_. Lysates were pelleted at 21,000 *g* for 10 min at 4 ºC, and 850 μL of supernatant were incubated with pre-washed anti-FLAG M2 magnetic beads (Sigma) beads for 1 hour at 4 ºC with end-over-end rotation. Beads were then washed 3 times with lysis buffer containing 350 mM NaCl. 1x Laemmli sample buffer (Bio-Rad) supplemented with 2-mercaptoethanol was added to beads directly to elute proteins. Samples were boiled at 95 ºC and analyzed by 12% SDS-PAGE and transferred to a 0.45 µm PVDF membrane. Anti-FLAG and anti-HA antibodies (Cell Signaling Technology) were used at a final concentration of 1:1000, and SuperSignal West Femto Maximum Sensitivity Substrate (ThermoFisher) was used to develop the blots. Blots were imaged by a ChemiDoc Imaging system (Bio-Rad). Images shown are representatives of 3 independent biological replicates.

### Incorporation assays

For co-producing CapRel^SJ46^ and Gp57, the SECΦ27 major capsid protein, single colonies of *E. coli* MG1655 containing pBR322-*capRel*^*SJ46*^ and pBAD33-*gp57* (wild-type or L114P variant) or corresponding empty vectors were grown overnight in M9-glucose. Overnight cultures were back-diluted to OD_600_ = 0.05 in 25 mL M9-glucose and grown to OD_600_ ∼ 0.3 at 37 ºC. Cells were pelleted at 4000 *g* for 5 min at 4 ºC and washed once with M9 (no glucose), and then back-diluted to OD_600_ = 0.1 in 15 mL M9 (no glucose) and recovered for 45 min at 37 ºC. At the beginning of the experiment, cells were induced with 0.2% arabinose. At the indicated time points (0, 10, 20, 30, 40 min), OD_600_ was measured and an aliquot of 250 µL of cells was transferred to microcentrifuge tube containing [5,6-^3^H]-uridine (PerkinElmer) (4 μCi/mL) for transcription measurements or EasyTag™ EXPRESS-^35^S Protein Labeling Mix, [^35^S] (PerkinElmer) at 44 μCi/mL for translation measurements. Tubes were incubated at 37 °C for 2 min, then quenched by addition of nonradioactive uridine (1.5 mM) or cysteine and methionine (15 mM each) and incubated for an additional 2 min. Samples were then added to ice cold trichloroacetic acid (TCA) (10% w/v) and incubated at least 30 min on ice to allow for precipitation. Resulting samples were vacuum filtered onto a glass microfiber filter (Whatman, 1820-024) that had been pre-wetted with 5% w/v TCA. Filters were washed with 35x volume of 5% w/v TCA, then with 5x volume of 100% ethanol. Air dried filters were placed in tubes with scintillation fluid and measured in a scintillation counter (PerkinElmer). CPM (Counts Per Million) was normalized to OD_600_ and percent incorporation at each time point was calculated by normalizing to T = 0. Data reported are the mean and individual data points from three independent biological replicates.

For producing the CapRel^SJ46^ N-terminal toxin domain, single colonies of *E. coli* MG1655 containing pBAD33-*capRel*^*SJ46*^*(1-272)* or an empty vector were grown overnight in M9-glucose. Transcription and translation experiments were done as described above. Data reported are the mean and individual data points from three independent biological replicates.

For phage infection experiments, single colonies of *E. coli* MG1655 harboring pBR322-EV or pBR322-*capRel*^*SJ46*^ were grown overnight in LB. Overnight cultures were back-diluted to OD_600_ = 0.05 in 25 mL fresh LB and grown to OD_600_ ∼ 0.3 at 25 ºC. Cells were then diluted to OD_600_ = 0.1 in 10 mL LB and infected with wild-type or mutant (L114P in Gp57, evolved clone 1 from population 3) SECΦ27 at MOI = 100 and incubated at 25 ºC. At the indicated time points (0, 15, 30, 45, 60 min), OD_600_ was measured and an aliquot of 250 µL of cells was transferred to a microcentrifuge tube containing [5,6-^3^H]-uridine (PerkinElmer) (32 μCi/mL) for transcription measurements or EasyTag™ EXPRESS-^35^S Protein Labeling Mix, [^35^S] (PerkinElmer) at 88 μCi/mL for translation measurements. Tubes were incubated at 25 °C for 4 min, then quenched by addition of nonradioactive uridine (1.5 mM) or cysteine and methionine (15 mM) and incubated for an additional 2 min. Samples were then processed same as above. Data reported are the mean and individual data points from three independent biological replicates. Statistical significance was determined by unpaired, two-tailed Student’s t-test (p<0.05).

### Homology search, alignment, and conservation analysis

CapRel^SJ46^ was identified in the sequence database from our previous bioinformatic survey of RSH proteins^24^ that included gene neighborhood analysis to identify TA systems^43^. Bacterial strains containing CapRel^SJ46^, CapRel^Ebc^ or CapRel^Kp^ with 100% amino acid identity were found on NCBI database. Local genomic regions (+/- 10kb of CapRel) were extracted and annotated for all coding sequences. Prophage genes and intact prophage regions were identified by PHASTER^44^. Additional homologs of CapRel^SJ46^ were identified by ConSurf^45^ using PSI-BLAST (default settings) to search UniRef90 database, yielding 44 homologs. For Fig S1, sequences were aligned with MAFFT L-INS-i v7.453 (Ref^46^) with manual curation of the C terminal region guided by homology modeling of the stand-alone Phrann Gp30 antitoxin using Swiss-Model^47^, and with our CapRel^SJ46^ predicted structure as a template. For Fig. S2, 52 homologs were used to generate the multiple sequence alignment by MAFFT and used as input for ConSurf. Conservation scores were calculated using the Bayesian method and default settings. An alignment of representative diverse sequences is shown and color-coded by percent identity (Fig. S2a).

Homologs of the major capsid proteins in BASEL phages were identified by BLASTp^48^ searches against each phage genome. Homologs of Gp57 (Gp8^Bas4^, Gp8^Bas5^, Gp8^Bas8^) were aligned by MUSCLE^49^ and colored by percent identity (Fig. S5a).

### CapRel^SJ46^ preparation for crystallization and HDX-MS

For the production of His_10_-SUMO-tagged CapRel^SJ46^ and CapRel^SJ46^ variants, *E. coli* BL21 (DE3) cells were transformed with pET24d plasmids containing the gene of interest and grown in LB medium to OD_600_ of 0.6. Expression of the protein of interest was induced by addition of 0.5 mM IPTG, and cells were grown for 3 hours at 30 °C. The culture was then centrifuged, and pellet was re-suspended in resuspension buffer (50 mM Tris-HCl pH 8.0, 1.5 M KCl, 2 mM MgCl_2_, 1 mM TCEP, 0,002% mellitic acid and 1 pastil of protease inhibitors cocktail (Roche)).

Cells were disrupted using a high-pressure homogenizer (Emulsiflex) and the supernatant was separated from the pellet by centrifugation and filtered through 0.45 μm filters. Protein extracts were loaded onto a gravity-flow column (Cytiva) packed with HisPur™ Nickel resin (ThermoFisher Scientific), washed with buffer A (50 mM Tris-HCl pH 8, 500 mM NaCl, 500 mM KCl, 1 mM TCEP, 0.002% Melitic acid) and stepwise eluted in the resuspension buffer supplemented with 500 mM imidazole. To remove remaining contaminants and imidazole, the elution fraction was immediately transferred to a size exclusion chromatography (SEC) column Superdex 200 PG column (GE Healthcare), previously equilibrated in the SEC buffer [50 mM HEPES pH 7.5, 500 mM NaCl, 500 mM KCl, 2 mM MgCl_2_, 1 mM TCEP, 0.002% mellitic acid (and 1 mM MnCl_2_ for all CapRel^SJ46^ proteins)]. The fractions containing the protein were concentrated to around 1 mg/mL and the His-tag was removed by incubating with UlpI protease (1:50 molar ratio) at 4 °C for 30 minutes. His_10_-SUMO-tag and the protease were then removed by passing the samples over a gravity-flow column (Cytiva) packed with HisPur™ Nickel resin (ThermoFisher Scientific). Purity of the sample preparation was assessed spectrophotometrically and by SDS-PAGE. For all the purified protein samples, OD_260_/OD_280_ ratio was below 0.6. Samples were stored at −20 °C or concentrated to 7 mg/ml and used directly in crystallization experiments.

For the purification of the His_10_-SUMO-CapRel^SJ46^ + His_10_-SUMO-Gp57 complex *E. coli* BL21 (DE3) strain containing freshly transformed pET24d-*His*_*10*_*-SUMO-capRel*^*SJ46*^*(Y155A)* and pET21a-*His*_*10*_*-SUMO-gp57* were grown in LB medium to OD_600_ of 0.2. This culture was then diluted in fresh LB media and grown until OD_600_ of 0.6. Expression of the protein of interest was induced by addition of 0.5 mM IPTG, and cells were grown for overnight at 16 °C. The subsequent purification, Sumo tag cleavage and purity assessment steps were identical to the workflow described above for the all the CapRel^SJ46^ protein variants.

### Crystallization of CapRel^SJ46^

The screening of crystallization conditions of CapRel^SJ46^ was carried out using the sitting-drop vapor-diffusion method. The drops were set up in Swiss (MRC) 96-well two-drop UVP sitting-drop plates using the Mosquito HTS system (TTP Labtech). Drops of 0.1 μL protein and 0.1 μL precipitant solution were equilibrated to 80 μL precipitant solution in the reservoir. Commercially available screens LMB and SG1 (Molecular Dimensions) were used to test crystallization conditions. The condition resulting in protein crystals (LMB screen position C9 for CapRel^SJ46^) were repeated as 2 µL drops. Crystals were harvested using suitable cryo-protecting solutions and vitrified in liquid N_2_ for transport and storage before X-ray exposure. X-ray diffraction data was collected at the SOLEIL synchrotron (Gif-sur-Yvette, Paris, France) on the Proxima 1 (PX1) and Proxima 2A (PX2A) beamlines using an Eiger-X 16M detector. Because of the high anisotropic nature of the data from all the crystals we performed anisotropic cutoff and correction of the merged intensity data as implemented on the STARANISO server (http://staraniso.globalphasing.org/) using the DEBYE and STARANISO programs. The analysis of the data suggested a resolution of 2.31 Å (with 2.31 Å in *a*\*, 2.85 Å in *b*\* and 2.72 Å in *c*\*).

### Structure determination

The data were processed with the XDS suite^50^ and scaled with Aimless. In all cases, the unit-cell content was estimated with the program MATTHEW COEF from the CCP4 program suite^51^. Molecular replacement was performed with Phaser^52^. The crystals of CapRel^SJ46^ diffracted on average to ≈2.3 Å. We used the coordinates of Rel_*Tt*_^NTD^ as search model for the toxSYNTH domain (PDBID 6S2T)^53^. The MR solution from Phaser was used in combination with Rosetta as implemented in the MR-Rosetta^54^ suit from the Phenix package^55^. After several iterations of manual building with Coot^56^ and maximum likelihood refinement as implemented in Buster/TNT^57^, the model was extended to cover all the residues (R/R_free_ of 21.5/26.0 %). **Table S1** details all the X-ray data collection and refinement statistics.

### Isothermal titration calorimetry (ITC)

For all ITC measurements CapRel^SJ46^ samples were prepared from the pET24d-*His*_*10*_*-SUMO-capRel*^*SJ46*^ as detailed above. In the case of Gp57, *E. coli* BL21 (DE3) cells were transformed with pET21a-*His*_*10*_*-SUMO-gp57* and grown in LB medium to OD_600_ of 0.2. This culture was then diluted in fresh LB media and grown until OD_600_ of 0.6. Expression of His_10_-SUMO*-*Gp57 was induced by addition of 0.1 mM IPTG, and cells were grown for overnight at 16 °C. The subsequent purification, SUMO-tag cleavage and purity assessment steps were identical to the workflow described above for the all the CapRel^SJ46^ protein variants. After removing the SUMO-tag, samples were concentrated to 10 μM and used directly for ITC immediately after purification.

All titrations were performed with an Affinity ITC (TA instruments) at 25 °C. For the titration, CapRel^SJ46^ was loaded in the instrument syringe at 150 μM and Gp57 was used in the cell at 10 μM. The titration was performed in 50 mM HEPES pH 7.5; 500 mM KCl; 500 mM; NaCl; 10 mM MgCl_2_; 1 mM TCEP; 0.002 % mellitic acid. Final concentrations were verified by the absorption using a Nanodrop One (ThermoScientific). All ITC measurements were performed by titrating 2 µL of CapRel^SJ46^ into Gp57 using a constant stirring rate of 75 rpm. All data were processed, buffer-corrected and analysed using the NanoAnalyse and Origin software packages.

### Hydrogen deuterium exchange mass spectrometry (HDX-MS)

Hydrogen Deuterium exchange mass spectrometry (HDX-MS) experiments were performed on an HDX platform composed of a Synapt G2-Si mass spectrometer (Waters Corporation) connected to a nanoAcquity UPLC system. Samples of CapRel^SJ46^ and CapRel^SJ46^ complexed with Gp57 were prepared at a concentration of 20 to 50 µM. For each experiment 5 µL of sample (CapRel^SJ46^ or CapRel^SJ46^-Gp57) were incubated for 1 min, 5 min, 15 min or 60 min in 95 µL of Labeling buffer L (50 mM HEPES, 500 mM KCl, 500 mM NaCl, 2 mM MgCl_2_, 1 mM TCEP, 0.002% mellitic acid, pH 7.5) at 20°C. The non-deuterated reference points were prepared by replacing buffer L by Equilibration buffer E (50 mM HEPES, 500 mM KCl, 500 mM NaCl, 2 mM MgCl_2_, 1 mM TCEP, 0.002% mellitic acid, pH 7.5). After labeling, the samples are quenched by mixing with 100 µL of pre-chilled quench buffer Q (1.2 % formic acid, pH 2.4). 70 µL of the quenched samples are directly transferred to the Enzymate BEH Pepsin Column (Waters Corporation) at 200 µL/min and at 20°C with a pressure 8.5 kPSI. Peptic peptides were trapped for 3 min on an Acquity UPLC BEH C18 VanGuard Pre-column (Waters Corporation) at a 200 µL/min flow rate in water (0.1% formic acid in HPLC water pH 2.5) before eluted to an Acquity UPLC BEH C18 Column for chromatographic separation. Separation was done with a linear gradient buffer (7-40% gradient of 0.1% formic acid in acetonitrile) at a flow rate of 40 µL/min. Peptides identification and deuteration upatke analysis was performed on the Synapt G2Si in ESI+ - HDMS^E^ mode (Waters Corporation). Leucine Enkephalin was applied for mass accuracy correction and sodium iodide was used as calibration for the mass spectrometer. HDMS^E^ data were collected by a 20-30 V transfer collision energy ramp. The pepsin column was washed between injections using pepsin wash buffer (1.5 M Gu-HCl, 4% (v/v) MeOH, 0.8% (v/v) formic acid). A blank run was performed between each sample to prevent significant peptide carry-over. Optimized peptide identification and peptide coverage for all samples was performed from undeuterated controls (five replicates). All deuterium time points were performed in triplicate.

### Data treatment and statistical analysis of HDX-MS

The non-deuterated references points were analyzed by PLGS (ProteinLynx Global Server 2.5.1, Waters) to identify the peptic peptides belonging CapRel^SJ46^ or Gp57. Then, all the HDMS^E^ data including reference and deuterated samples were processed by DynamX 3.0 (Waters) for deuterium uptake determination. We chose the following filtering parameters: minimum intensity of 1000, minimum and maximum peptide sequence length of 5 and 20, respectively, minimum MS/MS products of 3, minimum products per amino acid of 0.27, minimum score of 5, and a maximum MH+ error threshold of 15 p.p.m. Data were analyzed at peptidic and overall level and manually curated by visual inspection of individual spectra. The overall level is based on the relative fractional uptake (RFU) that can be calculated by the following formula:

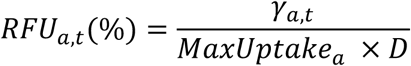

where *γ*_*a, t*_ is the deuterium uptake for peptide a at incubation time t, and *MaxUptake*_*a*_ *× D* is the theorical maximum uptake in deuterium value that peptide a can take. The ΔRFU compared RFU value between two different experiments conditions, in this case, this is the comparison between CapRel^SJ46^ and CapRel^SJ46^ + Gp57. Heat maps have been generated in DynamX. All the raw data can be accessed at: doi.org/10.6084/m9.figshare.19745089.

### CapRel^SJ46^ expression and purification for biochemical assays

Full-length *capRel*^*SJ46*^ was overexpressed in freshly transformed *E. coli* BL21(DE3) pET24d-*N-His*_*10*_*-SUMO-capRel*^*SJ46*^ pMG25-*paSpo* (VH-4) co-transformed with the plasmid encoding PaSpo Small Alarmone Hydrolase (SAH) from *Salmonella* phage SSU5 that had been shown to neutralize the toxicity of other toxSAS toxins^24^. Fresh transformants were used to inoculate 800 mL of LB medium (final OD_600_ of 0.03) supplemented with 50 μg/mL kanamycin, 20 μg/mL chloramphenicol and 0.2% arabinose. Bacterial cultures were grown at 37 °C until an OD_600_ of 0.4-0.5 and protein expression was induced with 0.1 mM IPTG (final concentration). Cells were grown for additional 1 hour at 30 °C and the biomass was harvested by centrifugation (10,000 rpm, for 5 minutes, JLA-10.500 rotor (Beckman Coulter)).

Cell mass was resuspended in buffer A (750 mM KCl, 500 mM NaCl, 5 mM MgCl_2_, 40 μM MnCl_2_, 40 μM Zn(OAc)_2_, 1 mM mellitic acid, 20 mM imidazole, 10% glycerol, 4 mM β-mercaptoethanol and 25 mM HEPES:KOH pH = 8) supplemented with 0.1 mM PMSF and 1 U/mL of DNase I, and lysed by one passage through a high-pressure cell disrupter (Stansted Fluid Power, 150 MPa). Mellitic acid was added to buffers as it was earlier shown to stabilise *Thermus thermophilus* Rel stringent factor^58^. Cell debris was removed by centrifugation (25,000 rpm for 1 hour, JA-25.50 rotor (Beckman Coulter)), the clarified lysate was filtered through a 0.22 μm syringe filter and loaded onto a HisTrap 5 ml HP column (Cytiva) pre-equilibrated in buffer A. The column was washed with 5 column volumes (CV) of buffer A, and the protein was eluted using a combination of stepwise and linear gradient (5 CV with 0-100% buffer B) of buffer B (750 mM KCl, 500 mM NaCl, 5 mM MgCl_2_, 40 μM MnCl_2_, 40 μM Zn(OAc)_2_, 1 mM mellitic acid, 1 M imidazole, 10% glycerol, 4 mM β-mercaptoethanol, 25 mM HEPES:KOH pH = 8). Fractions enriched in CapRel^SJ46^ (approximately 40% buffer B) were pooled, totalling approximately 5 mL. The sample was loaded on a HiLoad 16/600 Superdex 200 PG column pre-equilibrated with a high-salt buffer (buffer C; 2 M NaCl, 5 mM MgCl_2_, 10% glycerol, 4 mM β-mercaptoethanol, 25 mM HEPES:KOH pH = 8). The fractions containing CapRel^SJ46^ were pooled and applied on a HiPrep 10/26 desalting column (GE Healthcare) pre-equilibrated with storage buffer (buffer D; 720 mM KCl, 5 mM MgCl_2_, 40 mM arginine, 40 mM glutamic acid, 10% glycerol, 4 mM β-mercaptoethanol, 25 mM HEPES:KOH pH = 8). Fractions containing CapRel^SJ46^ were collected (about 14 mL in total) and the His_10_-SUMO tag was cleaved off by addition of 10 µg of His_6_-Ulp1 per 1 mg CapRel^SJ46^ followed by a 30-minute incubation on ice. After the His_10_-SUMO tag was cleaved off, the protein was passed through a 5 mL HisTrap HP pre-equilibrated with buffer D supplemented with 20 mM imidazole. Fractions containing CapRel^SJ46^ in the flow-through were collected and concentrated on an Amicon Ultra (Millipore) centrifugal filter device with a 10 kDa cut-off. The purity of protein preparations was assessed by SDS-PAGE. Protein preparations were aliquoted, frozen in liquid nitrogen and stored at –80 °C. Individual single-use aliquots were discarded after the experiment.

### Cell-free translation

Experiments with PURExpress *in vitro* protein synthesis kit (NEB, E6800) were performed as per the manufacturer’s instructions. All reactions were supplemented with 0.8 U/µL RNase Inhibitor Murine (NEB, M0314S). Purified CapRel^SJ46^ protein was used at a final concentration of 250 nM, with *gp57* or *gp57*(*L114P I115F)* as template plasmid at 10 ng/µL. As a mock control CapRel^SJ46^ was substituted for equal volume of HEPES:Polymix buffer, pH = 7.5. After a 10-minute incubation at 37 °C, a 1.34 µL aliquot of the reaction mixture was taken and quenched by addition of 13.66 µL of 2x sample buffer (100 mM Tris:HCl pH = 6.8, 4% SDS, 0.02% bromophenol blue, 20% glycerol, 20 mM DTT and 4% β-mercaptoethanol), and DHFR template plasmid was added to the remaining reaction mixture at a final concentration of 20 ng/µL. After further incubation at 37 °C for 1 hour, the reaction mixture was mixed with 9-fold volume of 2x sample buffer and 5 µL of the mixture was resolved on 18% SDS-PAGE gel. The SDS-PAGE gel was fixed by incubating for 5 min at room temperature in 50% ethanol solution supplemented with 2% phosphoric acid, washed three times with water for 20 min at room temperature, and stained with “blue silver” solution (0.12% Brilliant Blue G250 (Sigma-Aldrich, 27815), 10% ammonium sulfate, 10% phosphoric acid, and 20% methanol) overnight at room temperature. After washing with water for 3 hours at room temperature, the gel was imaged on an Amersham™ ImageQuant 800 (Cytiva) imaging system. For tRNA pyrophosphorylation experiments (see below), Gp57 and Gp57(L114P I115F) were produced in similar reaction mixture without CapRel^SJ46^ and DHFR template at 37 °C for 2 hours.

### tRNA pyrophosphorylation by CapRel^J46^

The reaction mixture containing 5 µM tRNA from *E. coli* MRE600 (Sigma-Aldrich, 10109541001), 500 µM γ^32^P-ATP, 250 nM CapRel^SJ46^ and 1/10 volume of either wild-type Gp57 or Gp57(L114P I115F) products from PUREsystem in HEPES:Polymix buffer, pH = 7.5 (5 mM Mg^2+^ final concentration) supplemented with 1 mM DTT was incubated at 37 °C for 10 min. To visualize phosphorylated tRNA, the reaction sample was mixed in 2 volumes of RNA dye (98% formamide, 10 mM EDTA, 0.3% bromophenol blue and 0.3% xylene cyanol), tRNA was denatured at 37 °C for 10 min and resolved on urea-PAGE in 1x TBE (8 M urea, 8% PAGE). The gel was stained with SYBR Gold (Life technologies, S11494) and exposed to an imaging plate overnight. The imaging plate was imaged by a FLA-3000 (Fujifilm).

---

**Figure S1.**
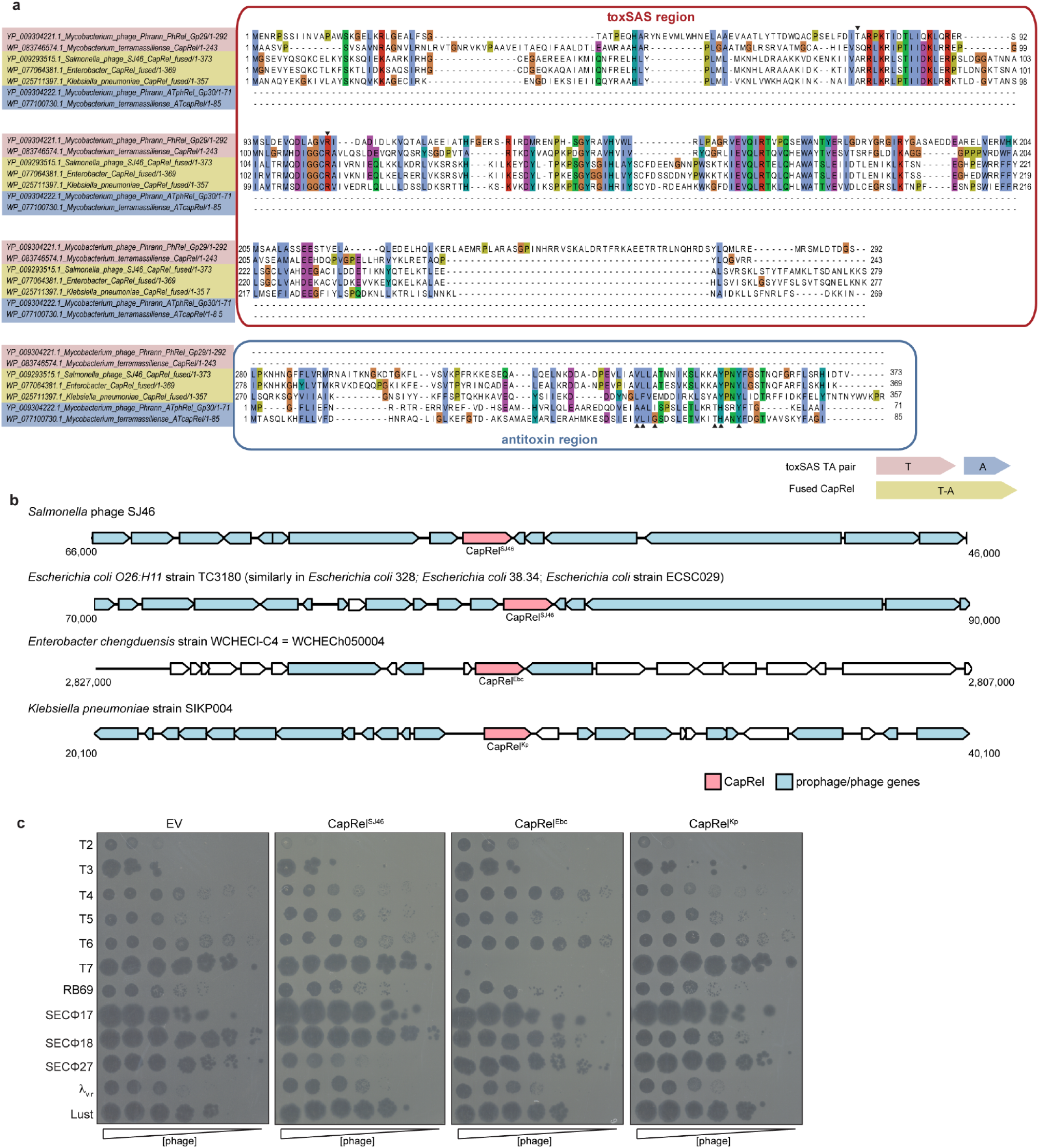
Analysis of CapRel homologs. (**a**) Sequence alignment comparing fused CapRel systems with related, unfused systems. Alignment of toxSAS PhRel and ATphRel from the *Mycobacterium* phage Phrann, non-fused CapRel and ATcapRel from *Mycobacterium terramassilience*, and the three fused systems CapRel^SJ46^, CapRel^Ebc^ and CapRel^Kp^. The N-terminal region of fused CapRel systems is a toxSAS toxin domain, while the C-terminal region is homologous to the antitoxins of the PhRel and unfused CapRel TA systems. Substituted sites of CapRel^SJ46^ (see Fig. 2g) are indicated with black arrowheads. The inset diagram summarises the homologous regions of the bicistronic toxin-antitoxin and fused toxin-antitoxin systems considered here. (**b**) Genome maps of native locations of CapRel^SJ46^, CapRel^Ebc^ and CapRel^Kp^ (+/- 10kb) with predicted flanking prophage and phage genes. (**c**) Serial dilutions of the phages indicated spotted on lawns of cells producing CapRel^SJ46^, CapRel^Ebc^, or CapRel^Kp^ or harboring an empty vector (EV).

**Figure S2.**
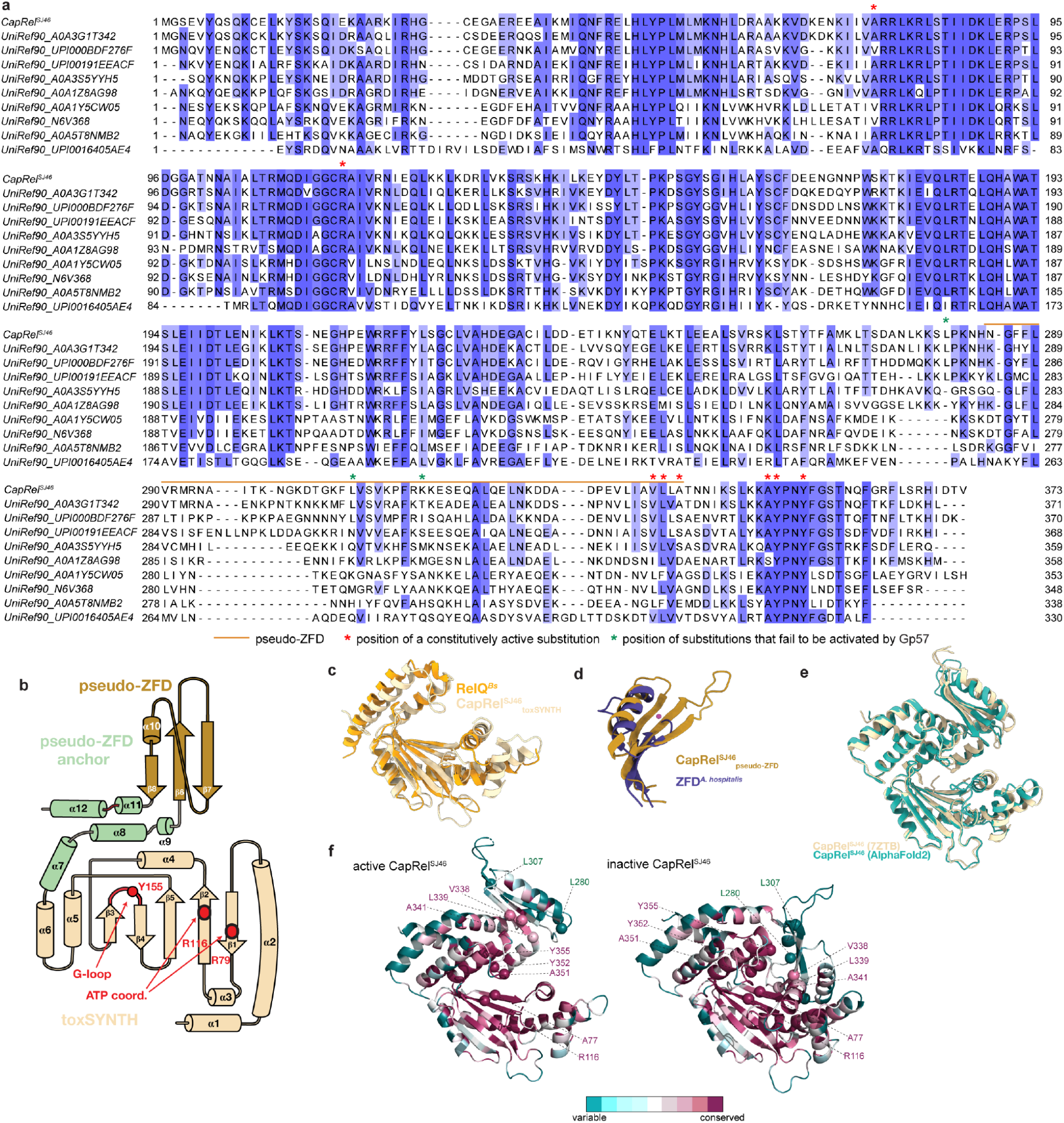
Structural analysis of CapRel^SJ46^. (**a**) Alignment of CapRel^SJ46^ and diverse fused CapRel homologs, with labels indicating that pseudo-ZFD and location of substitutions that render CapRel^SJ46^ constitutively active or unable to be activated by Gp57, the SECΦ27 major capsid protein. (**b**) Topology of CapRel^SJ46^. The toxSYNTH domain is colored in light yellow, the pseudo-ZFD in dark gold and the regions that anchor pseudo-ZFD to toxSYNTH are in green. The adenine coordinating R79 and R116 are shown as red dots and the G-loop is colored in red. (**c**) Superposition of the toxSYNTH domain of CapRel^SJ46^ (colored in light yellow) onto RelQ (PDBID: 5DEC, colored in light orange) from *Bacillus subtilis*. (**d)** Superposition of the pseudo-ZFD of CapRel^SJ46^ (colored in dark gold) onto the ZFD transcription factor of *Acidianus hospitalis* (2LVH, colored in purple). (**e**) Superposition of the crystal structure of CapRel^SJ46^ (colored in light yellow) onto the structure of the open state predicted by AlphaFold (colored in green). (**f)** Structures of the open (*left*; from crystal structure) or closed (*right*; AlphaFold prediction) conformations of CapRel^SJ46^ color coded by the conservation score of each amino acid calculated by ConSurf. Substitutions that render CapRel^SJ46^ constitutively active mutants or unable to be activated by Gp57 are labeled as spheres.

**Figure S3.**
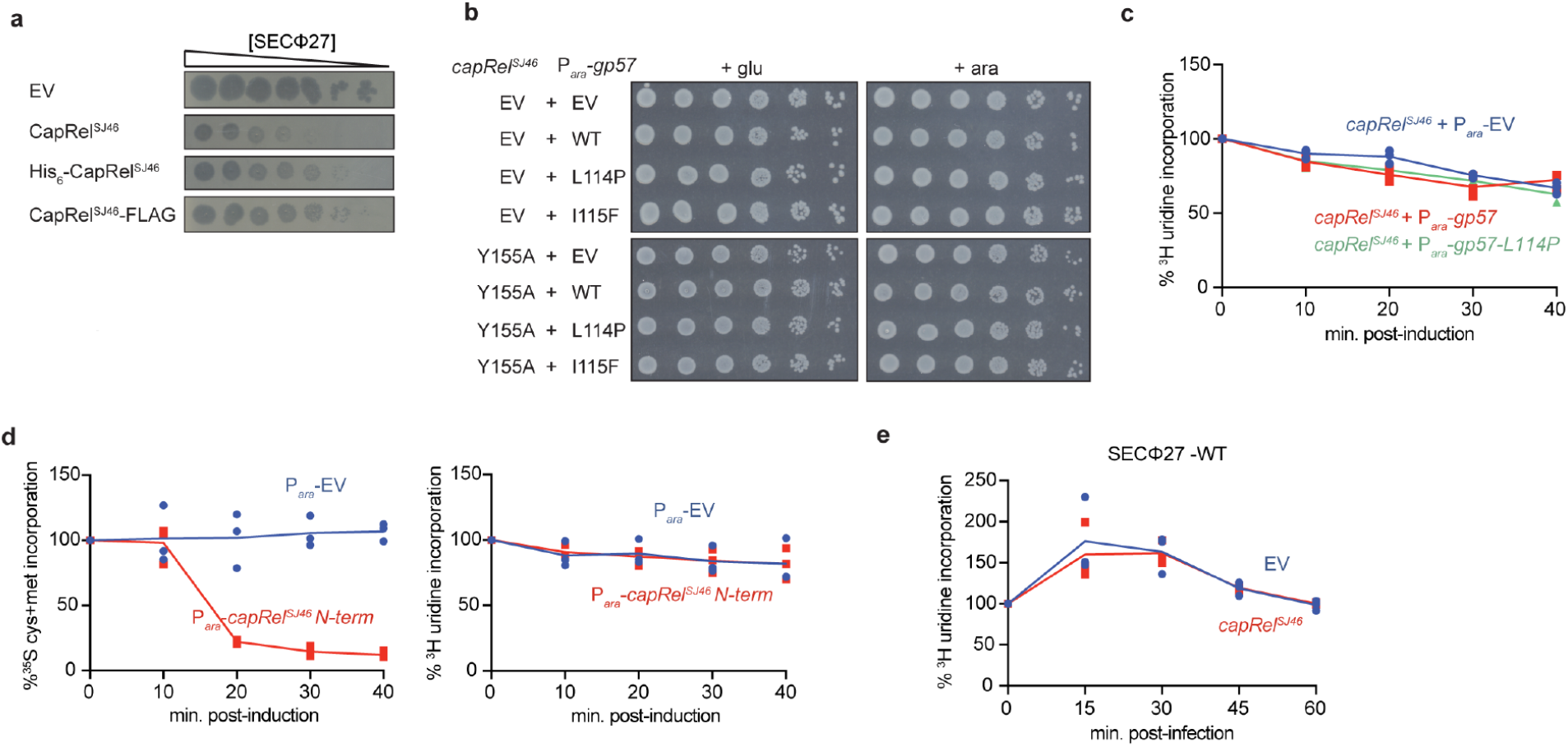
Gp57 from SECΦ27 triggers CapRel^SJ46^ to inhibit translation, not transcription. (**a**) Serial dilutions of phage SECΦ27 spotted on lawns of cells producing CapRel^SJ46^, His_6_-CapRel^SJ46^, or CapRel^SJ46^-FLAG, or harboring an empty vector (EV). (**b**) Serial dilutions on media containing glucose (*left*) or arabinose (*right*) of cells expressing *capRel*^*SJ46*^*(Y155A)* from its native promoter or an empty vector and expressing the indicated variant of Gp57 from an arabinose-inducible promoter. (**c**) Cells harboring CapRel^SJ46^ and producing the wild-type or L114P mutant of Gp57 (expressed from an arabinose-inducible promoter) or harboring an empty vector were pulse-labeled with ^3^H-uridine at the times indicated post-addition of arabinose. (**d**) Cells producing the CapRel^SJ46^ N-terminal toxin domain (expressed from an arabinose-inducible promoter) or harboring an empty vector were pulse-labeled with ^35^S-Cys/Met (*left*) or ^3^H-uridine (*right*) at the times indicated post-addition of arabinose. (**e**) Same as (**d**) but for cells carrying CapRel^SJ46^ or an empty vector and at times post-infection with SECΦ27 at MOI = 100.

**Figure S4.**
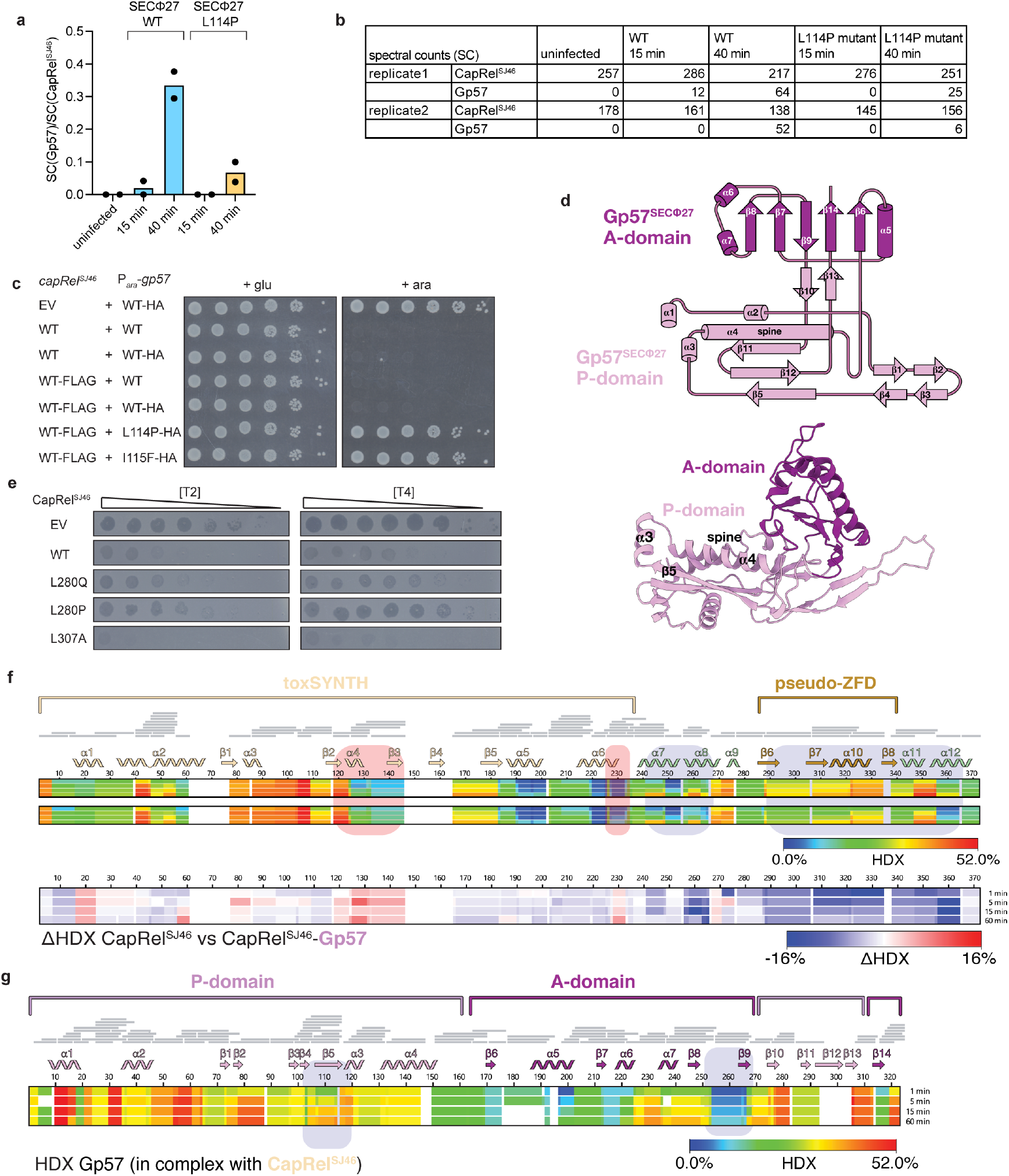
Characterization of the CapRel^SJ46^-Gp57 interaction. (**a**) Immunoprecipitation of CapRel^SJ46^-FLAG from cells infected with wild-type SECΦ27 or mutant phage that produces Gp57(L114P), followed by mass spectrometry. Spectrum counts (SC) of Gp57 that had co-precipitated with CapRel^SJ46^ were normalized to the spectrum counts of CapRel^SJ46^. (**b**) Same as in (**a**) but showing spectrum counts of CapRel^SJ46^ and Gp57 in two independent replicates. (**c**) Serial dilutions on media containing glucose (*left*) or arabinose (*right*) of cells producing CapRel^SJ46^ or CapRel^SJ46^-FLAG, each expressed from its native promoter, and the indicated variant of untagged or HA-tagged version of Gp57, expressed from an arabinose-inducible promoter. (**d**) Topology and cartoon representation of SECΦ27 Gp57. The P-domain is colored in pink and the A-domain in violet. (**e**) Serial dilutions of T2 and T4 phage spotted on cells producing the indicated mutant of CapRel^SJ46^ or harboring an empty vector. (**f**) Heat maps representing the HDX of CapRel^SJ46^ (*top*) and CapRel^SJ46^-Gp57 complex (*center*) and the “HDX (*bottom*). Regions involved in strong uptake such as residues 115-145 and 225-235 (which includes the active site #-strand #2 and the G-loop) are shaded in red and regions involved in strong protection 240-268 and 288-366 (which include both anchors and the pseudo-ZFD) are shaded in blue. (**g**) Heat map representing the HDX of Gp57 in the complex with CapRel^SJ46^. Shaded regions highlight areas of variable HDX signal that indicate these regions are involved in the CapRel^SJ46^-Gp57 interface.

**Figure S5.**
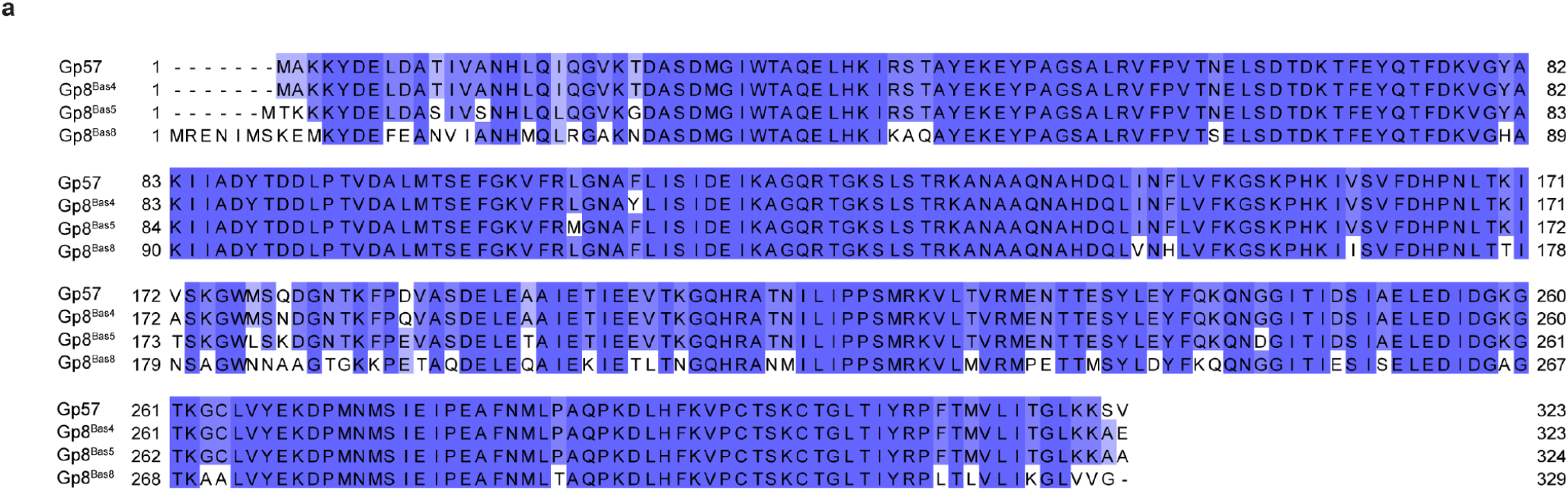
The major capsid protein from multiple, related phages activate CapRel^SJ46^. (**a**) Multiple sequence alignment of the major capsid proteins from phages SECΦ27, Bas4, Bas5 and Bas8.

