## Supplemental Tables for "Direct activation of an innate immune system in bacteria by a viral capsid protein"

**Table S1. X-ray data collection and processing.**
The $CC_{1/2}$ criterion was used to determine the resolution range. Values for the outer shell are given in parentheses.

| **Sample** | **CapRel$^{SJ46}$** |
|---|---|
| Diffraction source | Soleil PX1 |
| Wavelength (Å) | 0.9786 |
| Temperature (K) | 100.0 |
| Detector | Eiger-X 16M |
| Crystal-detector distance (mm) | 332.44 |
| Rotation range per image (°) | 0.01 |
| Exposure time per image (s) | 0.01 |
| Space group | $P2_1$ |
| *a*, *b*, *c* (Å) | 49.6 136.2 57.8 |
| α, β, γ (°) | 90.0, 102.7, 90.0 |
| Mosaicity (°) | 0.20 |
| Resolution range (Å) | 68.10 – 2.31 |
| Total N°. of reflections | 147846 (7291) |
| N°. of unique reflections | 20877 (1044) |
| Completeness (ellipsoidal %) | 91.9 (63.1) |
| Redundancy | 7.1 (7.0) |
| $\langle I/\sigma(I) \rangle$ | 11.2 (1.5) |
| $CC_{1/2}$ | 0.998 (0.712) |
| $R_{pim}$ | 0.051 (0.531) |
| Overall *B* factor / Wilson plot (Å$^2$) | 54.7 |
| R-factor (%) | 21.2 |
| $R_{free}$-factor (%) | 26.5 |
| Ramachandran profile (%) | |
| Core | 97.9 |
| Allowed | 2.1 |
| Outliers | 0.0 |
| R.m.s. deviations | |
| Bond lengths (Å) | 0.013 |
| Bond angles (°) | 1.52 |
| Number of atoms | 5534 |
| Macromolecules | 5304 |
| Solvent | 228 |
| Other | 3 |
| B-factors (Å$^2$) | |
| All atoms | 68.7 |
| Macromolecules | 69.2 |
| Solvent atoms | 55.8 |
| Other atoms | 63.6 |
| PDB ID | 7ZTB |

**Table S2. Mass spectrometry analysis of SECΦ27 phage lysates (wild type and mutant producing Gp57(L114P)).**

| | | | WT Gp57 | | | Gp57(L114P) mutant | | |
|---|---|---|---|---|---|---|---|---|
| Gene product | annotation | MW (kDa) | Spectrum count (SC) | SC/MW | coverage (%) | Spectrum count (SC) | SC/MW | coverage (%) |
| Gp57 | DUF2184 domain-containing protein | 36 | 878 | 24.39 | 86 | 548 | 15.22 | 92 |
| Gp55 | HtjA | 18 | 325 | 18.06 | 88 | 190 | 10.56 | 87 |
| Gp63 | tail protein/depolymerase | 25 | 333 | 13.32 | 79 | 220 | 8.80 | 78 |
| Gp66 | tail tape-measure protein | 98 | 197 | 2.01 | 58 | 168 | 1.71 | 56 |
| Gp81 | tail fiber protein | 72 | 121 | 1.68 | 44 | 88 | 1.22 | 44 |
| Gp52 | portal protein | 48 | 105 | 2.19 | 71 | 102 | 2.13 | 71 |
| Gp71 | tail fiber protein | 132 | 99 | 0.75 | 39 | 73 | 0.55 | 35 |
| Gp56 | scaffolding protein | 28 | 68 | 2.43 | 67 | 69 | 2.46 | 69 |
| Gp77 | recombinase | 24 | 40 | 1.67 | 65 | 15 | 0.63 | 36 |
| Gp20 | hypothetical protein | 16 | 11 | 0.69 | 43 | 13 | 0.81 | 43 |
| Gp78 | ssDNA-binding protein | 17 | 24 | 1.41 | 77 | 16 | 0.94 | 66 |
| Gp59 | head-tail adaptor | 16 | 11 | 0.69 | 26 | 10 | 0.63 | 17 |
| Gp62 | structural protein | 15 | 20 | 1.33 | 44 | 16 | 1.07 | 44 |
| Gp51 | terminase large subunit | 59 | 10 | 0.17 | 11 | 5 | 0.08 | 8.4 |
| Gp54 | major head subunit precursor | 41 | 6 | 0.15 | 6.2 | 15 | 0.37 | 26 |
| Gp68 | minor tail protein L | 28 | 15 | 0.54 | 33 | 9 | 0.32 | 21 |
| Gp60 | ribonucleoside-triphosphate reductase | 14 | 10 | 0.71 | 29 | 9 | 0.64 | 29 |
| Gp1 | DNA adenine methyltransferase | 28 | 4 | 0.14 | 12 | 6 | 0.21 | 26 |
| Gp79 | chaperone of endosialidase | 26 | 14 | 0.54 | 41 | 9 | 0.35 | 33 |
| Gp14 | DNA-cytosine methylase | 27 | 9 | 0.33 | 18 | 9 | 0.33 | 20 |
| Gp67 | minor tail protein | 13 | 13 | 1.00 | 49 | 8 | 0.62 | 26 |
| Gp75 | exodeoxyribonuclease | 41 | 5 | 0.12 | 12 | 7 | 0.17 | 18 |
| Gp39 | ATP-binding protein | 23 | 5 | 0.22 | 20 | 5 | 0.22 | 20 |
| Gp70 | tail assembly protein | 21 | 6 | 0.29 | 35 | 5 | 0.24 | 31 |
| Gp72 | hypothetical protein | 22 | 4 | 0.18 | 16 | 0 | 0.00 | 0 |
| Gp3 | hypothetical protein | 7 | 5 | 0.71 | 19 | 4 | 0.57 | 19 |
| Gp53 | head morphogenesis protein | 29 | 5 | 0.17 | 11 | 2 | 0.07 | 8 |
| Gp73 | outer membrane protein | 10 | 4 | 0.40 | 42 | 0 | 0.00 | 0 |
| Gp80 | hypothetical protein | 11 | 3 | 0.27 | 21 | 2 | 0.18 | 21 |
| Gp22 | hypothetical protein | 10 | 0 | 0.00 | 0 | 4 | 0.40 | 24 |
| Gp69 | C40 family peptidase | 28 | 3 | 0.11 | 9.5 | 0 | 0.00 | 0 |

**Table S3. Strains**

**Bacterial Strains**

| Name | Genotype | Source |
|---|---|---|
| ML6 | MG1655 | |
| | DH5α | Invitrogen |
| ML3836 | BW27783 | |
| | BL21 (DE3) | Lab stock |
| ML3837 | MG1655 pBAD33-EV pEXT20-EV | this study |
| ML3838 | MG1655 pBAD33-*capRel*$^{SJ46}$*(1-272)* pEXT20-EV | this study |
| ML3839 | MG1655 pBAD33-*capRel*$^{SJ46}$*(1-272)* pEXT20-*capRel*$^{SJ46}$*(273-373)* | this study |
| ML3840 | MG1655 pBR322-EV | this study |
| ML3841 | MG1655 pBR322-*capRel*$^{SJ46}$ | this study |
| ML3842 | MG1655 pBR322-*capRel*$^{Ebc}$ | this study |
| ML3843 | MG1655 pBR322-*capRel*$^{Kp}$ | this study |
| ML3844 | MG1655 pBR322-*capRel*$^{SJ46}$*(Y155A)* | this study |
| ML3845 | MG1655 pBR322-*capRel*$^{Ebc}$*(Y153A)* | this study |
| ML3846 | MG1655 pBR322-*capRel-chimera* | this study |
| ML3847 | MG1655 pBAD33-*capRel*$^{SJ46}$ | this study |
| ML3848 | MG1655 pBAD33-*capRel*$^{SJ46}$*(A77K)* | this study |
| ML3849 | MG1655 pBAD33-*capRel*$^{SJ46}$*(R116A)* | this study |
| ML3850 | MG1655 pBAD33-*capRel*$^{SJ46}$*(V338A)* | this study |
| ML3851 | MG1655 pBAD33-*capRel*$^{SJ46}$*(L339A)* | this study |
| ML3852 | MG1655 pBAD33-*capRel*$^{SJ46}$*(A341K)* | this study |
| ML3853 | MG1655 pBAD33-*capRel*$^{SJ46}$*(A351K)* | this study |
| ML3854 | MG1655 pBAD33-*capRel*$^{SJ46}$*(Y352A)* | this study |
| ML3855 | MG1655 pBAD33-*capRel*$^{SJ46}$*(Y355A)* | this study |
| ML3856 | MG1655 pBR322-*His*$_6$-*capRel*$^{SJ46}$ | this study |
| ML3857 | MG1655 pBR322-*capRel*$^{SJ46}$ pBAD33-EV | this study |
| ML3858 | MG1655 pBR322-*capRel*$^{SJ46}$ pBAD33-*gp57* | this study |
| ML3859 | MG1655 pBR322-*capRel*$^{SJ46}$ pBAD33-*gp57(L114P)* | this study |
| ML3860 | MG1655 pBR322-*capRel*$^{SJ46}$ pBAD33-*gp57(I115F)* | this study |

| | | |
|---|---|---|
| ML3861 | MG1655 pBR322-*capRel*$^{SJ46}$ pBAD33-*gp57-HA* | this study |
| ML3862 | MG1655 pBR322-*capRel*$^{SJ46}$-*FLAG* pBAD33-*gp57-HA* | this study |
| ML3863 | MG1655 pBR322-*capRel*$^{SJ46}$-*FLAG* pBAD33-*gp57(L114P)-HA* | this study |
| ML3864 | MG1655 pBR322-*capRel*$^{SJ46}$-*FLAG* pBAD33-*gp57(I115F)-HA* | this study |
| ML3865 | MG1655 pBR322-EV pBAD33-*gp57* | this study |
| ML3866 | MG1655 pBR322-*capRel*$^{SJ46}$*(L280Q)* pBAD33-*gp57* | this study |
| ML3867 | MG1655 pBR322-*capRel*$^{SJ46}$*(L280P)* pBAD33-*gp57* | this study |
| ML3868 | MG1655 pBR322-*capRel*$^{SJ46}$*(L307A)* pBAD33-*gp57* | this study |
| ML3869 | MG1655 pBR322-*capRel*$^{SJ46}$*(L280Q)* | this study |
| ML3870 | MG1655 pBR322-*capRel*$^{SJ46}$*(L280P)* | this study |
| ML3871 | MG1655 pBR322-*capRel*$^{SJ46}$*(L307A)* | this study |
| ML3872 | MG1655 pBR322-*capRel*$^{SJ46}$*(L280Q)-FLAG* pBAD33-*gp57-HA* | this study |
| ML3873 | MG1655 pBR322-*capRel*$^{SJ46}$*(L280P)-FLAG* pBAD33-*gp57-HA* | this study |
| ML3874 | MG1655 pBR322-*capRel*$^{SJ46}$*(L307A)-FLAG* pBAD33-*gp57-HA* | this study |
| ML3875 | MG1655 pBR322-*capRel*$^{SJ46}$ pBAD33-*gp8*$^{Bas4}$ | this study |
| ML3876 | MG1655 pBR322-*capRel*$^{SJ46}$ pBAD33-*gp8*$^{Bas5}$ | this study |
| ML3877 | MG1655 pBR322-*capRel*$^{SJ46}$ pBAD33-*gp8*$^{Bas8}$ | this study |
| ML3878 | MG1655 pBR322-*capRel*$^{SJ46}$ pBAD33-*gp8*$^{Bas4}$*(Y113F)* | this study |
| ML3879 | MG1655 pBR322-*capRel*$^{SJ46}$ pBAD33-*gp57(F113Y)* | this study |
| ML3880 | MG1655 pBR322-*capRel*$^{SJ46}$-*FLAG* | this study |
| ML3881 | MG1655 pBR322-EV pBAD33-EV | this study |
| ML3882 | MG1655 pBR322-EV pBAD33-*gp57* | this study |
| ML3883 | MG1655 pBR322-EV pBAD33-*gp57(L114P)* | this study |
| ML3884 | MG1655 pBR322-EV pBAD33-*gp57(I115F)* | this study |
| ML3885 | MG1655 pBR322-*capRel*$^{SJ46}$*(Y155A)* pBAD33-EV | this study |
| ML3886 | MG1655 pBR322-*capRel*$^{SJ46}$*(Y155A)* pBAD33-*gp57* | this study |
| ML3887 | MG1655 pBR322-*capRel*$^{SJ46}$*(Y155A)* pBAD33-*gp57(L114P)* | this study |
| ML3888 | MG1655 pBR322-*capRel*$^{SJ46}$*(Y155A)* pBAD33-*gp57(I115F)* | this study |
| ML3889 | MG1655 pBAD33-EV | this study |
| ML3890 | MG1655 pBAD33-*capRel*$^{SJ46}$*(1-272)* | this study |
| ML3891 | MG1655 pBR322-EV pBAD33-*gp57-HA* | this study |
| ML3892 | MG1655 pBR322-*capRel*$^{SJ46}$-*FLAG* pBAD33-*gp57* | this study |

| | | |
|---|---|---|
| ML3893 | BW27783 pBR322-EV pBAD33-*gp8*$^{Bas8}$ | this study |
| ML3894 | BW27783 pBR322-*capRel*$^{SJ46}$ pBAD33-*gp8*$^{Bas8}$ | this study |
| ML3895 | BW27783 pBR322-*capRel*$^{SJ46}$ pBAD33-*gp8*$^{Bas8}$*(F120L)* | this study |
| ML3896 | BW27783 pBR322-*capRel*$^{SJ46}$ pBAD33-*gp8*$^{Bas8}$*(I124F)* | this study |
| ML3897 | DH5α pBAD33-EV | this study |
| ML3898 | DH5α pBAD33-*capRel*$^{SJ46}$*(1-272)* | this study |
| ML3899 | DH5α pEXT20-EV | this study |
| ML3900 | DH5α pEXT20-*capRel*$^{SJ46}$*(273-373)* | this study |
| ML3901 | DH5α pBR322-EV | this study |
| ML3902 | DH5α pBR322-*capRel*$^{SJ46}$ | this study |
| ML3903 | DH5α pBR322-*capRel*$^{SJ46}$*(Y155A)* | this study |
| ML3904 | DH5α pBR322-*capRel*$^{Ebc}$ | this study |
| ML3905 | DH5α pBR322-*capRel*$^{Ebc}$*(Y153A)* | this study |
| ML3906 | DH5α pBR322-*capRel*$^{Kp}$ | this study |
| ML3907 | DH5α pBR322-*capRel-chimera* | this study |
| ML3908 | DH5α pBAD33-*capRel*$^{SJ46}$ | this study |
| ML3909 | DH5α pBAD33-*capRel*$^{SJ46}$*(A77K)* | this study |
| ML3910 | DH5α pBAD33-*capRel*$^{SJ46}$*(R116A)* | this study |
| ML3911 | DH5α pBAD33-*capRel*$^{SJ46}$*(V338A)* | this study |
| ML3912 | DH5α pBAD33-*capRel*$^{SJ46}$*(L339A)* | this study |
| ML3913 | DH5α pBAD33-*capRel*$^{SJ46}$*(A341K)* | this study |
| ML3914 | DH5α pBAD33-*capRel*$^{SJ46}$*(A351K)* | this study |
| ML3915 | DH5α pBAD33-*capRel*$^{SJ46}$*(Y352A)* | this study |
| ML3916 | DH5α pBAD33-*capRel*$^{SJ46}$*(Y355A)* | this study |
| ML3917 | DH5α pBR322-*His*$_6$*-capRel*$^{SJ46}$ | this study |
| ML3918 | DH5α pBAD33-*gp57* | this study |
| ML3919 | DH5α pBAD33-*gp57(L114P)* | this study |
| ML3920 | DH5α pBAD33-*gp57(I115F)* | this study |
| ML3921 | DH5α pBR322-*capRel*$^{SJ46}$*-FLAG* | this study |
| ML3922 | DH5α pBAD33-*gp57-HA* | this study |
| ML3923 | DH5α pBAD33-*gp57(L114P)-HA* | this study |
| ML3924 | DH5α pBAD33-*gp57(I115F)-HA* | this study |

| | | |
|---|---|---|
| ML3925 | DH5α pBR322-*capRel*$^{SJ46}$*(L280Q)* | this study |
| ML3926 | DH5α pBR322-*capRel*$^{SJ46}$*(L280P)* | this study |
| ML3927 | DH5α pBR322-*capRel*$^{SJ46}$*(L307A)* | this study |
| ML3928 | DH5α pBR322-*capRel*$^{SJ46}$*(L280Q)-FLAG* | this study |
| ML3929 | DH5α pBR322-*capRel*$^{SJ46}$*(L280P)-FLAG* | this study |
| ML3930 | DH5α pBR322-*capRel*$^{SJ46}$*(L307A)-FLAG* | this study |
| ML3931 | DH5α pBAD33-*gp8*$^{Bas4}$ | this study |
| ML3932 | DH5α pBAD33-*gp8*$^{Bas5}$ | this study |
| ML3933 | DH5α pBAD33-*gp8*$^{Bas8}$ | this study |
| ML3934 | DH5α pBAD33-*gp8*$^{Bas4}$*(Y113F)* | this study |
| ML3935 | DH5α pBAD33-*gp57(F113Y)* | this study |
| ML3936 | DH5α pBAD33-*gp8*$^{Bas8}$*(F120L)* | this study |
| ML3937 | DH5α pBAD33-*gp8*$^{Bas8}$*(I124F)* | this study |
| ML3938 | MG1655 pBAD33-*capRel*$^{SJ46}$*(R78A)* | this study |
| ML3939 | MG1655 pBAD33-*capRel*$^{SJ46}$*(K311A)* | this study |
| ML3940 | MG1655 pBAD33-*capRel*$^{SJ46}$*(R314A)* | this study |
| ML3941 | MG1655 pBAD33-*capRel*$^{SJ46}$*(E319A)* | this study |
| ML3942 | MG1655 pBAD33-*capRel*$^{SJ46}$*(K346A)* | this study |
| ML3943 | DH5α pBAD33-*capRel*$^{SJ46}$*(R78A)* | this study |
| ML3944 | DH5α pBAD33-*capRel*$^{SJ46}$*(K311A)* | this study |
| ML3945 | DH5α pBAD33-*capRel*$^{SJ46}$*(R314A)* | this study |
| ML3946 | DH5α pBAD33-*capRel*$^{SJ46}$*(E319A)* | this study |
| ML3947 | DH5α pBAD33-*capRel*$^{SJ46}$*(K346A)* | this study |
| VH-1 | DH5α pET24d-*gp57* | this study |
| VH-2 | DH5α pET24d-*gp57(L114P I115F)* | this study |
| VH-3 | DH5α pET24d-*N-His*$_{10}$*-SUMO-capRel*$^{SJ46}$ | this study |
| VH-4 | BL21 (DE3) pET24d-*N-His*$_{10}$*-SUMO-capRel*$^{SJ46}$ pMG25-*paSpo* | this study |
| AGP-1 | BL21 (DE3) pET21a-*His*$_{10}$*-SUMO-gp57* | this study |
| AGP-2 | BL21 (DE3) pET24d-*N-His*$_{10}$*-SUMO-capRel*$^{SJ46}$ | this study |

**Phage Strains**

| Name | Genotype | Source |
|---|---|---|
| phML1 | T2 | ATCC Cat #: 11303-B2 |

| | | |
|---|---|---|
| phML2 | T3 | ATCC Cat #: 11303-B3 |
| phML3 | T4 | ATCC Cat #: 11303-B4 |
| phML4 | T5 | ATCC Cat #: 11303-B5 |
| phML5 | T6 | ATCC Cat #: 11303-B6 |
| phML6 | T7 | Gift from R. Sorek |
| phML7 | RB69 | Laval Collection, HER #158 |
| phML8 | $\lambda_{vir}$ | Gift from R. Sorek |
| phML9 | SECΦ17 | (Doron et al., 2018) |
| phML10 | SECΦ18 | (Doron et al., 2018) |
| phML11 | SECΦ27 | (Doron et al., 2018) |
| phML12 | Lust | (Malki et al., 2016) |
| phML43 | SECΦ27 evolved clone 1 from population 3 containing Gp57(L114P) | this study |
| phML44 | SECΦ27 evolved clone 1 from population 1 containing Gp57(I115F) | this study |
| phML45 | Bas4 | (Maffei et al., 2021) |
| phML46 | Bas5 | (Maffei et al., 2021) |
| phML47 | Bas8 | (Maffei et al., 2021) |
| phML48 | Bas8 escape clone 1 containing Gp8(F120L) | this study |
| phML49 | Bas8 escape clone 2 containing Gp8(I124F) | this study |
| phML50 | Bas4 clone 1 containing Gp8(Y113F) | this study |
| phML51 | Bas4 clone 2 containing Gp8(Y113F) | this study |

**Table S4. Plasmids**

| Plasmid | Description | Source |
|---|---|---|
| pBAD33-EV | empty vector of pBAD33 (p15A ori, $P_{ara}$ promoter) | lab stock |
| pBAD33-*capRel*$^{SJ46}$*(1-272)* | arabinose inducible N-terminal fragment (1-272) of CapRel$^{SJ46}$ | this study |
| pEXT20-EV | empty vector of pEXT20 (pBR322 ori, $P_{lac}$ promoter) | lab stock |
| pEXT20-*capRel*$^{SJ46}$*(273-373)* | IPTG inducible C-terminal fragment (273-373) of CapRel$^{SJ46}$ | this study |
| pBR322-EV | derivative of pBR322 with $P_{tet}$ removed | lab stock |
| pBR322-*capRel*$^{SJ46}$ | *capRel*$^{SJ46}$ with native promoter | this study |
| pBR322-*capRel*$^{SJ46}$*(Y155A)* | *capRel*$^{SJ46}$*(Y155A)* with native promoter | this study |
| pBR322-*capRel*$^{Ebc}$ | *capRel*$^{Ebc}$ with native promoter | this study |
| pBR322-*capRel*$^{Ebc}$*(Y153A)* | *capRel*$^{Ebc}$*(Y153A)* with native promoter | this study |
| pBR322-*capRel*$^{Kp}$ | *capRel*$^{Kp}$ with native promoter | this study |
| pBR322-*capRel-chimera* | *capRel* chimera producing CapRel$^{SJ46}$ (272-341) replaced with CapRel$^{Ebc}$ (270-339) | this study |
| pBAD33-*capRel*$^{SJ46}$ | arabinose inducible *capRel*$^{SJ46}$ | this study |
| pBAD33-*capRel*$^{SJ46}$*(A77K)* | arabinose inducible c*apRel*$^{SJ46}$*(A77K)* | this study |
| pBAD33-*capRel*$^{SJ46}$*(R116A)* | arabinose inducible *capRel*$^{SJ46}$*(R116A)* | this study |
| pBAD33-*capRel*$^{SJ46}$*(V338A)* | arabinose inducible *capRel*$^{SJ46}$*(V338A)* | this study |
| pBAD33-*capRel*$^{SJ46}$*(L339A)* | arabinose inducible *capRel*$^{SJ46}$*(L339A)* | this study |
| pBAD33-*capRel*$^{SJ46}$*(A341K)* | arabinose inducible *capRel*$^{SJ46}$*(A341K)* | this study |
| pBAD33-*capRel*$^{SJ46}$*(A351K)* | arabinose inducible *capRel*$^{SJ46}$*(A351K)* | this study |
| pBAD33-*capRel*$^{SJ46}$*(Y352A)* | arabinose inducible *capRel*$^{SJ46}$*(Y352A)* | this study |
| pBAD33-*capRel*$^{SJ46}$*(Y355A)* | arabinose inducible *capRel*$^{SJ46}$*(Y355A)* | this study |
| pBAD33-*capRel*$^{SJ46}$*(R78A)* | arabinose inducible *capRel*$^{SJ46}$*(R78A)* | this study |
| pBAD33-*capRel*$^{SJ46}$*(K311A)* | arabinose inducible *capRel*$^{SJ46}$*(K311A)* | this study |
| pBAD33-*capRel*$^{SJ46}$*(R314A)* | arabinose inducible *capRel*$^{SJ46}$*(R314A)* | this study |
| pBAD33-*capRel*$^{SJ46}$*(E319A)* | arabinose inducible *capRel*$^{SJ46}$*(E319A)* | this study |
| pBAD33-*capRel*$^{SJ46}$*(K346A)* | arabinose inducible *capRel*$^{SJ46}$*(K346A)* | this study |
| pBR322-*His*$_6$*-capRel*$^{SJ46}$ | *capRel*$^{SJ46}$ with N-terminal *His*$_6$*-tag* | this study |
| pBAD33-*gp57* | arabinose inducible *gp57* | this study |

| | | |
|---|---|---|
| pBAD33-*gp57(L114P)* | arabinose inducible *gp57(L114P)* | this study |
| pBAD33-*gp57(I115F)* | arabinose inducible *gp57(I115F)* | this study |
| pBR322-*capRel*$^{SJ46}$-*FLAG* | *capRel*$^{SJ46}$ with C-terminal *FLAG-tag* | this study |
| pBAD33-*gp57-HA* | arabinose inducible *gp57* with C-terminal *HA-tag* | this study |
| pBAD33-g*p57(L114P)-HA* | arabinose inducible *gp57(L114P)* with C-terminal *HA-tag* | this study |
| pBAD33-g*p57(I115F)-HA* | arabinose inducible *gp57(I115F)* with C-terminal *HA-tag* | this study |
| pBR322-*capRel*$^{SJ46}$*(L280Q)* | *capRel*$^{SJ46}$*(L280Q)* with native promoter | this study |
| pBR322-*capRel*$^{SJ46}$*(L280P)* | *capRel*$^{SJ46}$*(L280P)* with native promoter | this study |
| pBR322-*capRel*$^{SJ46}$*(L307A)* | *capRel*$^{SJ46}$*(L307A)* with native promoter | this study |
| pBR322-*capRel*$^{SJ46}$*(L280Q)-FLAG* | *capRel*$^{SJ46}$*(L280Q)* with C-terminal *FLAG-tag* | this study |
| pBR322-*capRel*$^{SJ46}$*(L280P)-FLAG* | *capRel*$^{SJ46}$*(L280P)* with C-terminal *FLAG-tag* | this study |
| pBR322-*capRel*$^{SJ46}$*(L307A)-FLAG* | *capRel*$^{SJ46}$*(L307A)* with C-terminal *FLAG-tag* | this study |
| pBAD33-*gp8*$^{Bas4}$ | arabinose inducible *gp8*$^{Bas4}$ | this study |
| pBAD33-*gp8*$^{Bas5}$ | arabinose inducible *gp8*$^{Bas5}$ | this study |
| pBAD33-*gp8*$^{Bas8}$ | arabinose inducible *gp8*$^{Bas8}$ | this study |
| pBAD33-*gp8*$^{Bas4}$*(Y113F)* | arabinose inducible *gp8*$^{Bas4}$*(Y113F)* | this study |
| pBAD33-*gp57(F113Y)* | arabinose inducible *gp57(F113Y)* | this study |
| pBAD33-*gp8*$^{Bas8}$*(F120L)* | arabinose inducible *gp8*$^{Bas8}$*(F120L)* | this study |
| pBAD33-*gp8*$^{Bas8}$*(I124F)* | arabinose inducible *gp8*$^{Bas8}$*(I124F)* | this study |
| pET24d-*gp57* | *gp57* expressed under the control of T7 promoter | this study |
| pET24d-g*p57(L114P I115F)* | *gp57(L114P I115F)* expressed under the control of T7 promoter | this study |
| pET24d-*His*$_{10}$-*SUMO* | *E. coli* codon-optimized *His*$_{10}$-*SUMO-tag* expressed under the control of T7 promoter | ref[59] |
| pET24d-*N-His*$_{10}$-*SUMO-capRel*$^{SJ46}$ | *capRel*$^{SJ46}$ with N-terminal *His*$_{10}$-*SUMO-tag* | this study |
| pMG25-*paSpo* | PaSpo SAH from *Salmonella* phage *SSU5* expressed under the control of IPTG inducible promoter $P_{A1/04/03}$ | ref[28] |

| | | |
|---|---|---|
| DHFR Control Template | DHFR expressed under the control of T7 promoter | NEB |
| pET21a-*His*$_{10}$-*SUMO-gp57* | *gp57* with *His*$_{10}$-*SUMO-tag* | this study |

**Table S5. Primers**

| Name | Purpose | Sequence (5'-3') |
|---|---|---|
| TZ-1 | *capRel*$^{SJ46}$ into pBR322 backbone (with native promoter) | GCGTATCACGAGGCCCTTTCGTCTTCAAGA<br>ATTCTCATGTTAGTTACGTTAGGTAACATTA<br>TGGGTATGTTTCATTTTCATACCCATTGATT<br>TCTAGCTTGATTTGTTATTATATGTAACAAC<br>ACATTAATCATTTGAACTGGATGTTAGTAA<br>TGGGCAGTGAAGTATATCAAAGCCAAAAAT<br>GCGAGCTTAAATATTCGAAATCTCAAATAG<br>AAAAAGCAGCCAGAAAAATTAGACATGGT<br>TGTGAGGGCGCAGAGCGAGAAGAAGCAAT<br>TAAAATGATTCAAAATTTCCGCGAGCTTCA<br>TTTATATCCATTGATGCTAATGAAAAATCA<br>TCTTGATAGGGCTGCAAAAAAAGTTGATAA<br>AGAGAATAAGATTATTGTTGCAAGACGCCT<br>AAAAAGACTTAGCACAATAATTGATAAACT<br>GGAGCGTCCAAGTCTTGACGGCGGCGCAAC<br>AAATAATGCAATAGCCCTTACTAGAATGCA<br>GGATATTGGCGGATGTAGAGCTATAGTTAG<br>AAACATCGAACAACTAAAGAAACTAAAAG<br>ACCGACTAGTTAAAAGTCGCTCCAAGCATA<br>AAATCTTAAAGGAATATGACTATTTAACTC<br>CAAAGCCAAGCGGATATAGTGGCATTCACC<br>TTGCATATAGTTGTTTTGATGAAGAAAACG<br>GTAACAATCCGTGGAGTAAAACTAAAATCG<br>AAGTACAACTAAGAACAGAACTCCAACAT<br>GCTTGGGCAACAAGTCTGGAGATAATAGAT<br>ACACTTGAAAATATTAAGCTGAAAACTTCT<br>AATGAAGGTCACCCTGAGTGGAGAAGATTT<br>TTTTATCTTTCTGGATGCCTAGTTGCTCATG<br>ATGAAGGTGCCTGTATTCTTGATGATGAAA<br>CCATCAAAAATTATCAAACAGAGCTTAAAA<br>CATTGGAGGAGGCTTTATCTGTTCGCTCAA<br>AATTAAGTACATATACATTTGCAATGAAGC<br>TGACATCAGATGCAAACCTAAAGAAAAGC<br>CTGCCCAAAAACCATAATGGATTCTTTTTG<br>GTGAGGATGAGAAACGCCATAACAAAAAA<br>TGGCAAGGATACTGGAAAGTTTCTTGTTTC<br>TGTGAAACCTTTTAGGAAAAAAGAATCGGA<br>ACAAGCCCTCCAGGAACTTAATAAGGATGA<br>TGCGGATCCTGAAGTATTAATTGCGGTGCT<br>GCTTGCCACTAACAACATTAAATCTCTTAA<br>AAAGGCTTATCCTAACTATTTTGGGTCAAC<br>AAATCAGTTCGGGAGATTCCTTAGCAGACA<br>CATTGATACTGTGTAAATTCCTATGCCTGGC<br>TTAGCCAGGCATAGCCAAAGATTGGGCACC |

| | | |
|---|---|---|
| | | TCGCTAACGGATTCACCACTCCAAGAATTGGAGC |
| TZ-2 | *capRel*$^{Ebc}$ into pBR322 backbone (with native promoter) -fragment 1 | CCTTTCGTCTTCAAGAATTCTCATGTTTACCCTACATACATCATTATTGACGGTATCAAGATCAGCCGCTCCCGCTTGTCATTTGCCTTTCAGTTGGAGTTAGCAAAAGCCCTTAAAATTTATGAAGAAAATCATACGTAAGTAGCATAAATGACTTGAGTTTTCAGCCAGTATGCATTCTCATGCGTACATACATGGCAATAATATTAACGGGAGGATATCGGGAATGGGCAATGAAGTTTATGAAAGCCAGAAGTGTACGCTTAAATTTAGTAAGACCCTTATTGATAAGTCCGCGCAGTCGATCCGTCATGGGTGTGATGGTGAGCAAAAAGCACAGGCTATTGCTATGATCCAGAACTTCCGTGAACTTCACCTTTATCCGTTGATGTTAATGAAAAATCACTTAGCCCGTGCCGCGGCAAAGATTGACAAAAAGACTATTGTCGCACGCCGCCTTAAGCGTTTAAGTACCATCATTGATAAATTGGAACGCCCCACCTTAGACAATGGAAAAACCAGCAATGCAATCCGTGTAACACGCATGCAGGATATTGGGGGTTGTCGTGCAATCGTTAAAAACATCGAACAGCTTAAAGAATTGCGCGAGCGCCTGGTTAAATCCCGCAGCGTACATAAAATCATCAAAGAGAGTGATTATTTGAC |
| TZ-3 | *capRel*$^{Ebc}$ into pBR322 backbone (with native promoter) -fragment 2 | AGAGAGTGATTATTTGACTCCAAAAGAGTCGGGGTACGGTGGTATCCACTTAGTATATAGCTGTTTTGATTCCAGTGATGACAATTACCCATGGAAAAAGACGAAGATCGAGGTACAACTTCGTACGCAGTTGCAGCACGCATGGGCAACATCTTTAGAGATCATCGATACGTTAGAGAACATTAAGTTGAAAACCTCTAGCGAGGGGCACGAAGATTGGCGCCGCTTTTTTATTTGAGCGGGTGTCTGGTAGCTCACGATGAGAAGGCCTGCGTGCTGGACAAGGAGGTAGTAAAGGAATACCAAAAGGAGTTGAAAGCCTTAGCGGAGCATCTGTCTGTCATTAGCAAGTTGGGTAGTTATGTGTTTTCACTTAGCGTGACTTCACAAAACGAACTTAAAAACAAAATTCCAAAAAACCATAAGGGACATTATCTGGTCACTATGAAGCGCGTGAAGGACGAACAGCAGCCTGGGAAAATTAAGTTTGAGGTGTCCGTGACTCCGTACCGTATTAACCAAGCCGATGAGGCCTTAGAGGCATTGAAACGTGATGATGCAAATCCGGAAGTGCCCATCGCTGTGTTGCTTGCGACCGAATCCGTGAAATCACTTAAAAAGGCATAC |

| | | |
|---|---|---|
| | | CCAAATTACCTGGGCTCCACTAACCAGTTT GCACGCTTTTTAAGCAAGCATATTAAGTAA GGCACCTCGCTAACGGATTCACCAC |
| TZ-4 | *capRel*$^{Kp}$ into pBR322 backbone (with native promoter) -fragment 1 | CCTTTCGTCTTCAAGAATTCTCATGTCATCC TCTGTTACACGGCGAGTTCCACCATGGGCA CAGCTTACGTTTGATTTAATGATTTATTCCA GCCCCTTCATGCGAGGGGGTTGGGCTAAGT CAGGCAAAACGAGACATGTTGCACATTGTA GTGTGATTGCATTGTATCAAAAATATCAGG AGGTTATAGTGCCAGAACTCAAGCTATTAG ATGAAGGCAGTCACATGGCTAATGCACAAT ACGAGAAGGGGAAGATTGTTTAGCTTACT CGAAAAATCAAGTTAAGAAAGCGGGCGAA TGTATTCGTAAAGAGAACGGAGATATTGAA AAGTCAATTGAAATCATTCAGCAATATCGC GCAGCACATTTATATCCGTTAATGATCGTG AAAAATTTAGTTTGGAAACATGCACAGAAG ATCAACAAAAACGCGATCATTGCACGTCGC TTAAAACGTCTTCCTACAATCATTGATAAA TTGCGTCGCAAAACCCTGGACGGGGTGACG GCTAATTCAATCGCAGTCACTCGCATGTCG GATATCGGTGGATGTCGTGTAATCGTGGAA GACCGTTTGCAACTGCTGCTTCTGGATTCCT CCTTAGATAAAAGTCGTACCACGCATAAGT CAAAGGTGAAGGATTACATTAAGTCGCCGA AACCTACTGGGTACCGTG |
| TZ-5 | *capRel*$^{Kp}$ into pBR322 backbone (with native promoter) -fragment 2 | GCCGAAACCTACTGGGTACCGTGGGATTCA TCGCATCTATTCCTGCTATGACCGCGACGA AGCGCACAAATGGAAAGGGTTCGACATCG AGGTGCAGTTACGTACAAAGTTGCAACATC TTTGGGCAACTACAGTAGAAGTAGTAGACT TGTGCGAGGGCCGCAGCTTAAAGACCAACC CGTTCGAATCGAATCCTTCGTGGATTGAGT TCTTCCGCCTTATGTCAGAATTTATCGCCGA TGAGGAAGGCTTTATTTACCTTTCCCCCCAG GACAAGAATTTACTGAAGACTCGCTTGATC TCTTTGAATAATAAGCTGAATGCGATTGAC AAACTGTTGTCTTTTAACCGTCTTTTTTCTG ATAAAAAGATTAACCTGTCTCAACGTAAGT CCGGGTATGTAATCATTGCCATTAAAGGTA ATTCTATCTATTATAAGTTTTTCAGCCCAAC TCAAAAACACAAAGCCGTAGAACAATACTC GATTATTGAAAAGGACGATGACTATAATGG CTTGTTCGTCGAGATGGACGATATTCGCAA ACTGTCATATGCGTATCCCAACTACTTAATT GATACGCGTTTTTTCATTGACAAGTTTGAAT |

| | | |
|---|---|---|
| | | TATACACCAATACAAATTATTGGGTCAAGCCACGTTAAGGCACCTCGCTAACGGATTCACCAC |
| TZ-6 | linearize pBR322 | GGCACCTCGCTAACGGAT |
| TZ-7 | linearize pBR322 | ACATGAGAATTCTTGAAGACGAAAGG |
| TZ-8 | generate Y155A in *capRel*$^{SJ46}$ | GCAAGTGGCATTCACCTTGCATATAGTTG |
| TZ-9 | generate Y155A in *capRel*$^{SJ46}$ | TCCGCTTGGCTTTGGAGTTAA |
| TZ-10 | generate Y153A in *capRel*$^{Ebc}$ | GTCGGGGGCGGGTGGTATCCACTTAGTATATAGCT |
| TZ-11 | generate Y153A in *capRel*$^{Ebc}$ | CCACCCGCCCCCGACTCTTTTGGAGTCAA |
| TZ-12 | linearize pBAD33 | AAGCTTGGCTGTTTTGGC |
| TZ-13 | linearize pBAD33 | CTCGAATTCGCTAGCCCAAA |
| TZ-14 | *capRel*$^{SJ46}$*(1-272)* and *capRel*$^{SJ46}$ into pBAD33 | TACCCGTTTTTTTGGGCTAGCGAATTCGAGTGAACTGGATGTTAGTAATGGGCAGT |
| TZ-15 | *capRel*$^{SJ46}$*(1-272)* into pBAD33 | CTTCTCTCATCCGCCAAAACAGCCAAGCTTTTATGATGTCAGCTTCATTGCAAATGT |
| TZ-16 | linearize pEXT20 | CGGGGATCCTCTAGAGTCGAC |
| TZ-17 | linearize pEXT20 | GGTACCGAGCTCGAATTCTGTTTC |
| TZ-18 | *capRel*$^{SJ46}$*(273-373)* into pEXT20 | AATTCGAGCTCGGTACCGGGAGGATATCGGGAATGGATGCAAACCTAAAGAAAAGCCTGC |
| TZ-19 | *capRel*$^{SJ46}$*(273-373)* into pEXT20 | GCAGGTCGACTCTAGAGGATCCCCGTTACACAGTATCAATGTGTCTGCTAAGGA |
| TZ-20 | linearize pBR322-*capRel*$^{SJ46}$ to generate pBR322-*capRel-chimera* | GTGTTGCTTGCGACTAACAACATTAAATCTCTTAAAAAGGCT |
| TZ-21 | linearize pBR322-*capRel*$^{SJ46}$ to generate pBR322-*capRel-chimera* | TAAGTTCGTTTTGTGATGTCAGCTTCATTGCAAATGTATATGTACTTAATT |
| TZ-22 | insert *capRel*$^{Ebc}$(270-339) to generate pBR322-*capRel-chimera* | AAGCTGACATCACAAAACGAACTTAAAAACAAAATTCCAAAAAACC |
| TZ-23 | insert *capRel*$^{Ebc}$(270-339) to generate pBR322-*capRel-chimera* | GTGTTGCTTGCGACTAACAACATTAAATCTCTTAAAAAGGCT |
| TZ-24 | *capRel*$^{SJ46}$ into pBAD33 | CTTCTCTCATCCGCCAAAACAGCCAAGCTTTTACACAGTATCAATGTGTCTGCTAAG |
| TZ-25 | generate A77K in *capRel*$^{SJ46}$ | TATTGTTAAAAGACGCCTAAAAAGACTTAG |

| | | |
|---|---|---|
| TZ-26 | generate A77K in *capRel*$^{SJ46}$ | CGTCTTTTAACAATAATCTTATTCTCTTTATCA |
| TZ-27 | generate R116A in *capRel*$^{SJ46}$ | CGGATGTGCGGCTATAGTTAGAAACATCGAACAAC |
| TZ-28 | generate R116A in *capRel*$^{SJ46}$ | ATAGCCGCACATCCGCCAATATCCTGC |
| TZ-29 | generate V338A in *capRel*$^{SJ46}$ | AATTGCGGCGCTGCTTGCCACTAACAACA |
| TZ-30 | generate V338A in *capRel*$^{SJ46}$ | AGCAGCGCCGCAATTAATACTTCAGGATCCGC |
| TZ-31 | generate L339A in *capRel*$^{SJ46}$ | TGCGGTGGCGCTTGCCACTAACAACATTAAATCTC |
| TZ-32 | generate L339A in *capRel*$^{SJ46}$ | GCAAGCGCCACCGCAATTAATACTTCAGGATCC |
| TZ-33 | generate A341K in *capRel*$^{SJ46}$ | GCTGCTTAAAACTAACAACATTAAATCTCTTAAAA |
| TZ-34 | generate A341K in *capRel*$^{SJ46}$ | TTAGTTTTAAGCAGCACCGCAATTAA |
| TZ-35 | generate A351K in *capRel*$^{SJ46}$ | TAAAAAGAAATATCCTAACTATTTTGGGTCAAC |
| TZ-36 | generate A351K in *capRel*$^{SJ46}$ | GGATATTTCTTTTTAAGAGATTTAATGTTGTT |
| TZ-37 | generate Y352A in *capRel*$^{SJ46}$ | AAAGGCTGCGCCTAACTATTTTGGGTCAACAAATC |
| TZ-38 | generate Y352A in *capRel*$^{SJ46}$ | TTAGGCGCAGCCTTTTTAAGAGATTTAATGTTG |
| TZ-39 | generate Y355A in *capRel*$^{SJ46}$ | TCCTAACGCGTTTGGGTCAACAAATCAGTTCGGG |
| TZ-40 | generate Y355A in *capRel*$^{SJ46}$ | CCAAACGCGTTAGGATAAGCCTTTTTAAGAG |
| TZ-41 | construct pBR322-*His*$_6$-*capRel*$^{SJ46}$ | TCACCATCACCACGGCAGCAGCGGCATGGGCAGTGAAGTATATCAAAG |
| TZ-42 | construct pBR322-*His*$_6$-*capRel*$^{SJ46}$ | CCGTGGTGATGGTGATGATGCATTACTAACATCCAGTTCAAATGATT |
| TZ-43 | *gp57* into pBAD33 | TACCCGTTTTTTTGGGCTAGCGAATTCGAGATGGCTAAAAAATATGATGAACTAGATGC |
| TZ-44 | *gp57* into pBAD33 | TTCTCTCATCCGCCAAAACAGCCAAGCTTTTAAACTGACTTTTTCAAGCCAGTAATAAGC |
| TZ-45 | construct pBR322-*capRel*$^{SJ46}$-*FLAG* | GCGATTACAAGGATGACGATGACAAATAAATTCCTATGCCTGGCTTAGC |

| | | |
|---|---|---|
| TZ-46 | construct pBR322-*capRel*$^{SJ46}$-*FLAG* | CATCCTTGTAATCGCCGCTGCTGCCCACAGTATCAATGTGTCTGCTAAGG |
| TZ-47 | construct pBAD33-*gp57-HA* | CTACCCGTATGATGTGCCGGACTATGCATAAAAGCTTGGCTGTTTTGGCGG |
| TZ-48 | construct pBAD33-*gp57-HA* | ACATCATACGGGTAGCCGCTGCTGCCAACTGACTTTTTCAAGCC |
| TZ-49 | generate L280Q in *capRel*$^{SJ46}$ | GAAAAGCCAGCCCAAAAACCATAATGGATTC |
| TZ-50 | generate L280Q in *capRel*$^{SJ46}$ | TTGGGCTGGCTTTTCTTTAGGTTTGCATCTG |
| TZ-51 | generate L280P in *capRel*$^{SJ46}$ | GAAAAGCCCGCCCAAAAACCATAATGGATTC |
| TZ-52 | generate L280P in *capRel*$^{SJ46}$ | TTGGGCGGGCTTTTCTTTAGGTTTGCATCTG |
| TZ-53 | generate L307A in *capRel*$^{SJ46}$ | AAAGTTTGCGGTTTCTGTGAAACCTTTTAGGAAAA |
| TZ-54 | generate L307A in *capRel*$^{SJ46}$ | GAAACCGCAAACTTTCCAGTATCCTTGCCA |
| TZ-55 | *gp8*$^{Bas4}$ into pBAD33 | TACCCGTTTTTTTGGGCTAGCGAATTCGAGATGGCTAAAAAATATGATGAACTAGATGCT |
| TZ-56 | *gp8*$^{Bas4}$ into pBAD33 | CTTCTCTCATCCGCCAAAACAGCCAAGCTTTTACTCAGCTTTTTTCAAGCCG |
| TZ-57 | *gp8*$^{Bas5}$ into pBAD33 | TACCCGTTTTTTTGGGCTAGCGAATTCGAGATGACAAAGAAAAAATATGATGAGCTAGAC |
| TZ-58 | *gp8*$^{Bas5}$ into pBAD33 | CTTCTCTCATCCGCCAAAACAGCCAAGCTTTTACGCAGCCTTTTTCAAGCCA |
| TZ-59 | *gp8*$^{Bas8}$ into pBAD33 | CCCGTTTTTTTGGGCTAGCGAATTCGAGATGAGGGAAAATATTATGTCTAAAGAAATGAA |
| TZ-60 | *gp8*$^{Bas8}$ into pBAD33 | CTTCTCTCATCCGCCAAAACAGCCAAGCTTTCAGCCTACTACAAGACCTTTAATCA |
| TZ-61 | generate Y113F in *gp8*$^{Bas4}$ | TAACGCTTTTCTGATCTCCATTGATGAGATTAAAG |
| TZ-62 | generate Y113F in *gp8*$^{Bas4}$ | ATCAGAAAAGCGTTACCCAGGCGGAA |
| TZ-63 | generate F113Y in *gp57* | TAACGCTTATCTGATCTCCATTGATGAGATTAAAG |
| TZ-64 | generate F113Y in *gp57* | ATCAGATAAGCGTTACCCAGGCGGAA |

| | | |
|---|---|---|
| TZ-65 | *gp57* and *gp57(L114P I115F)* cloned into pET24d | CTTTAAGAAGGAGATATACCATGGCTAAAAAATATGATGAAC |
| TZ-66 | *gp57* and *gp57(L114P I115F)* cloned into pET24d | GCCGGATCTCAGTGGTGGTGTTAAACTGACTTTTTCAAGC |
| TZ-67 | *gp57* | CTGTGTGAAATTGTTATCCGAAAAATAAGGAGGAAAAAAAAATGGCTAAAAAATATGATGAACTAGATGCTACGATTGTAGCGAATCATTTGCAGATTCAGGGTGTCAAGACCGACGCTTCTGATATGGGTATTTGGACCGCTCAAGAGCTACACAAGATCCGCTCAACCGCATACGAGAAAGAATATCCGGCAGGTTCCGCGCTTCGCGTATTCCCTGTAACAAACGAGCTTTCTGATACTGATAAGACTTTTGAGTATCAGACTTTTGATAAGGTTGGCTACGCGAAAATTATCGCCGACTACACCGACGATCTGCCGACCGTGGACGCGCTGATGACTTCTGAATTTGGCAAGGTGTTCCGCCTGGGTAACGCTTTTCTGATCTCCATTGATGAGATTAAAGCAGGTCAGCGAACTGGCAAGAGCCTGTCAACTCGCAAGGCTAACGCCGCGCAAAATGCACATGATCAGCTGATTAACTTCCTGGTGTTCAAAGGTTCCAAGCCTCATAAGATCGTTTCCGTTTTCGATCATCCTAACCTTACGAAAATTGTTTCTAAAGGATGGATGAGCCAGGATGGAAACACCAAGTTCCCTGATGTGGCAAGCGATGAACTGGAGGCTGCAATCGAAACGATCGAGGAAGTAACCAAAGGTCAGCACCGAGCGACTAACATCCTGATCCCGCCGTCCATGCGCAAAGTCCTGACGGTTCGAATGGAAAATACCACTGAAAGTTATCTGGAATACTTCCAGAAGCAAAACGGCGGCATCACTATCGACTCTATCGCAGAGCTTGAGGATATTGACGGCAAAGGTACTAAAGGTTGCTTGGTTTACGAAAAAGATCCAATGAACATGAGCATTGAGATTCCAGAAGCGTTTAACATGCTTCCGGCGCAGCCAAAAGACCTTCATTTCAAGGTTCCTTGCACTTCCAAGTGTACTGGCCTTACGATTTACCGTCCGTTTACGATGGTGCTTATTACTGGCTTGAAAAAGTCAGTTTAAGCTTGGACTCCTGTTGATAG |
| TZ-68 | *gp57(L114P I115F)* | CTGTGTGAAATTGTTATCCGAAAAATAAGGAGGAAAAAAAAATGGCTAAAAAATATGATGAACTAGATGCTACGATTGTAGCGAATCATTTGCAGATTCAGGGTGTCAAGACCGACGCTTCTGATATGGGTATTTGGACCGCTCAAGAGCTACACAAGATCCGCTCAACCGCATACGAGAAAGAATATCCGGCAGGTTCCGCGCTTCGC |

| | | |
|---|---|---|
| | | GTATTCCCTGTAACAAACGAGCTTTCTGAT ACTGATAAGACTTTTGAGTATCAGACTTTT GATAAGGTTGGCTACGCGAAAATTATCGCC GACTACACCGACGATCTGCCGACCGTGGAC GCGCTGATGACTTCTGAATTTGGCAAGGTG TTCCGCCTGGGTAACGCTTTTCCGTTTTCCA TTGATGAGATTAAAGCAGGTCAGCGAACTG GCAAGAGCCTGTCAACTCGCAAGGCTAACG CCGCGCAAAATGCACATGATCAGCTGATTA ACTTCCTGGTGTTCAAAGGTTCCAAGCCTC ATAAGATCGTTTCCGTTTTCGATCATCCTAA CCTTACGAAAATTGTTTCTAAAGGATGGAT GAGCCAGGATGGAAACACCAAGTTCCCTGA TGTGGCAAGCGATGAACTGGAGGCTGCAAT CGAAACGATCGAGGAAGTAACCAAAGGTC AGCACCGAGCGACTAACATCCTGATCCCGC CGTCCATGCGCAAAGTCCTGACGGTTCGAA TGGAAAATACCACTGAAAGTTATCTGGAAT ACTTCCAGAAGCAAAACGGCGGCATCACTA TCGACTCTATCGCAGAGCTTGAGGATATTG ACGGCAAAGGTACTAAAGGTTGCTTGGTTT ACGAAAAAGATCCAATGAACATGAGCATT GAGATTCCAGAAGCGTTTAACATGCTTCCG GCGCAGCCAAAAGACCTTCATTTCAAGGTT CCTTGCACTTCCAAGTGTACTGGCCTTACG ATTTACCGTCCGTTTACGATGGTGCTTATTA CTGGCTTGAAAAAGTCAGTTTAAGCTTGGA CTCCTGTTGATAG |
| TZ-69 | Linearized pET24d plasmid without $His_6$-tag | CACCACCACTGAGATCCGGC |
| TZ-70 | Linearized pET24d plasmid without $His_6$-tag | GGTATATCTCCTTCTTAAAGTTAAACAAAA TTATTTC |
| TZ-71 | *capRel*$^{SJ46}$ cloned into pET24d-*N-His*$_{10}$*-SUMO* | ATCGCGAACAGATTGGTGGTGGCAGTGAAG TATATCAAAGCCAAAAATGCGAGC |
| TZ-72 | *capRel*$^{SJ46}$ cloned into pET24d-*N-His*$_{10}$*-SUMO* | GGTGGTGGTGGTGCTCGAGTTCTCATCCGC CAAAACAGCC |
| TZ-73 | Linearized pET24d-*N-His*$_{10}$*-SUMO* | ACCACCAATCTGTTCGCGATGAGC |
| TZ-74 | Linearized pET24d-*N-His*$_{10}$*-SUMO* | ACTCGAGCACCACCACCAC |
| TZ-75 | generate R78A in *capRel*$^{SJ46}$ | TGTTGCAGCGCGCCTAAAAAGACTTAGCAC A |

| | | |
|---|---|---|
| TZ-76 | generate R78A in *capRel*$^{SJ46}$ | AGGCGCGCTGCAACAATAATCTTATTCTCTTT |
| TZ-77 | generate K311A in *capRel*$^{SJ46}$ | TTCTGTGGCGCCTTTTAGGAAAAAAGAATCGGAAC |
| TZ-78 | generate K311A in *capRel*$^{SJ46}$ | AAAGGCGCCACAGAAACAAGAAACTTTCCAG |
| TZ-79 | generate R314A in *capRel*$^{SJ46}$ | ACCTTTTGCGAAAAAAGAATCGGAACAAGCCCTC |
| TZ-80 | generate R314A in *capRel*$^{SJ46}$ | TTTTTCGCAAAAGGTTTCACAGAAACAAGAAAC |
| TZ-81 | generate E319A in *capRel*$^{SJ46}$ | AGAATCGGCGCAAGCCCTCCAGGAACTTAATAAG |
| TZ-82 | generate E319A in *capRel*$^{SJ46}$ | GCTTGCGCCGATTCTTTTTTCCTAAAAGGTTTC |
| TZ-83 | generate K346A in *capRel*$^{SJ46}$ | CAACATTGCGTCTCTTAAAAAGGCTTATCCTAAC |
| TZ-84 | generate K346A in *capRel*$^{SJ46}$ | AGAGACGCAATGTTGTTAGTGGCAAGCAG |
